# The STA1–DOT2 interaction promotes nuclear speckle formation and splicing robustness in growth and heat stress responses

**DOI:** 10.64898/2026.01.11.698856

**Authors:** Heejin Kim, Kyoung-jae Yu, So Young Park, Dong Hye Seo, Woo Taek Kim, Dae-Jin Yun, Jae-Hoon Jung, Dong-Hoon Jeong, Byeong-ha Lee

## Abstract

Pre-mRNA splicing is carried out by the spliceosome, a large and dynamic ribonucleoprotein complex. The spliceosome is known to be stored in nuclear speckles (NS), which are now recognized as active subnuclear organelles for splicing. However, it remains poorly understood how spliceosomal protein-protein interactions are functionally coupled to NS organization to maintain splicing robustness in plants. Here, we report the functional significance of a specific interaction between two U4/U6·U5 tri-snRNP components of the spliceosome, STA1 and DOT2, in regulating NS organization, pre-mRNA splicing, and heat stress responses in *Arabidopsis*. We identified a missense mutation in *DOT2* (a Snu66/SART1 homolog) from a genetic suppressor of the PRP6 homolog mutant *sta1-1* (named *S307*). This mutation restored the weakened interaction between STA1 and DOT2 in the *sta1-1* mutant background. Genetic, biochemical, and cell biological analyses showed that variation in the strength of the STA1-DOT2 interaction was closely associated with changes in NS formation, splicing efficiency, as well as growth and heat tolerance. Pharmacological inhibition of STA1-associated NS formation by tubercidin recapitulated *sta1-1*-like phenotypes and splicing defects, supporting a functional link between NS organization and splicing outcomes. In addition, heat-induced weakening of the STA1-DOT2 interaction was accompanied by reduced NS formation and increased intron retention at the transcriptome-wide level including key heat-responsive transcripts. Based on these observations, we propose that the STA1-DOT2 interaction, likely reflecting the assembly state of the U4/U6·U5 tri-snRNP, functions as a heat-sensitive interaction node that couples spliceosome assembly to NS organization and splicing robustness under stress conditions.

## INTRODUCTION

Pre-mRNA splicing is an essential step in eukaryotic gene expression and is carried out by the spliceosome, a large and dynamic ribonucleoprotein complex composed of five small nuclear ribonucleoproteins (snRNPs) and numerous auxiliary factors (Albakri et al., 2025; Will and Lührmann, 2011). Among spliceosomal subcomplexes, the U4/U6·U5 tri-snRNP plays a pivotal role in spliceosome activation, and undergoes extensive structural rearrangements during the splicing cycle (Sander et al., 2006; Will and Lührmann, 2011). Defects in tri-snRNP assembly or stability result in splicing failure, underscoring the importance of precise and coordinated interactions among its components (Klimesová et al., 2021; Schneider et al., 2010).

*Arabidopsis* STABILIZED1 (STA1) is homologous to the pre-mRNA splicing factors PRP1, Prp6p, and PRPF6 (hereafter collectively referred to as PRP6, unless otherwise indicated) in *Schizosaccharomyces pombe*, *Saccharomyces cerevisiae*, and *Homo sapiens*, respectively. Within the tri-snRNP, PRP6 resides in the U5 snRNP and interacts with components of the U4/U6 di-snRNP, thereby physically bridging the U5 snRNP and the U4/U6 di-snRNP (Bertram et al., 2017; Liu et al., 2006; Makarov et al., 2000; Nguyen et al., 2016; Zhan et al., 2018). Consistent with this role, depletion or mutation of PRP6 in yeast and human cells disrupts tri-snRNP formation and leads to the accumulation of U4/U6 di-snRNP, indicating that PRP6 is essential for tri-snRNP assembly (Galisson and Legrain, 1993; Schaffert et al., 2004).

The *Arabidopsis sta1-1* mutant carries a 6-bp deletion within the half-a-tetratricopeptide (HAT) repeat domain of STA1 (Lee et al., 2006). HAT domains are known to mediate protein–protein interactions and promote the assembly of multiprotein complexes (Zeytuni and Zarivach, 2012). Supporting this, yeast two-hybrid assays using human tri-snRNP components have shown that PRPF6 interacts with several tri-snRNP components through its HAT domain (Liu *et al*., 2006). These findings suggest that the PRP6 homolog STA1 contributes to pre-mRNA splicing in plants primarily by promoting proper spliceosome assembly through its interactions with other tri-snRNP components.

In the nucleus, nuclear speckles (NS) are prominent subnuclear membraneless compartments enriched in spliceosomal proteins and RNAs (Ilik et al., 2020; Smith et al., 2020). NS serve as sites for the storage, assembly, and dynamic exchange of splicing factors and are closely associated with transcriptionally active chromatin, and are increasingly recognized as sites that actively contribute to efficient pre-mRNA processing (Daguenet et al., 2012; Wu et al., 2024). NS are thought to form through the self-assembly of resident proteins and RNAs and to be maintained by transient protein-protein and protein-RNA interactions (Choi et al., 2025; Day et al., 2012; Kim et al., 2016; Paul et al., 2024). Importantly, localization of spliceosomal components to NS enhances the splicing efficiency of nearby genes, whereas impaired speckle association leads to splicing defects (Liu et al., 2021; Wang et al., 2024).

Several lines of evidence suggest that tri-snRNP components contribute directly to NS organization. In human cells, depletion of PRPF6 disrupts the NS localization of another tri-snRNP component PRPF31 and causes splicing defects (Yildirim et al., 2021). In plants, NS localization of splicing factors has also been implicated in stress adaptation. For example, two alternatively spliced isoforms of RDM16 (a tri-snRNP component PRPF3 homolog) cooperatively promote nuclear condensation and enhance heat tolerance in *Arabidopsis* (Ma et al., 2025). These studies support the idea that NS organization is functionally linked to spliceosome integrity and splicing outcomes, particularly under stress. However, it remains unclear how the interaction-dependent assembly state of the tri-snRNP contributes to NS organization in plants.

In this study, from a *sta1-1* suppressor line (*S307*), we identified a missense mutation in *DEFECTIVELY ORGANIZED TRIBUTARIES2* (*DOT2*; also known as *MERISTEM-DEFECTIVE*, *MDF* (de Luxan-Hernandez et al., 2022; Petricka et al., 2008)), hereafter referred to as *dot2^S^*. DOT2 is a U4/U6·U5 tri-snRNP component homologous to yeast Snu66p and human SART1, which are known to bind to and stabilize the PRP6-involved bridge within the tri-snRNP (Bertram *et al*., 2017; Zhan *et al*., 2018). The *dot2^S^* suppressor allele restores the weakened interaction between STA1 and DOT2 in the *sta1-1* background, leading to recovery of NS formation, improved pre-mRNA splicing, and restoration of growth and heat tolerance. Our results support the idea that the STA1-DOT2 interaction functions as a critical and heat-sensitive interaction node within the tri-snRNP, contributing to spliceosome assembly to NS organization, and splicing robustness under stress conditions.

## RESULTS

### A mutation in DOT2, a U4/U6·U5 tri-snRNP-associated protein, suppresses the *sta1-1* defects

The *sta1-1* mutants, defective in the PRP6 homolog *STA1* gene, are small in size and hypersensitive to heat (Kim et al., 2017; Lee *et al*., 2006). To elucidate the molecular mechanism underlying these phenotypes, we employed a forward genetic approach to identify *sta1-1* modifiers, aiming to uncover genetic interactors of *STA1*. From an ethyl methanesulfonate (EMS)-treated *sta1-1* mutant pool, we isolated a *sta1-1* suppressor *S307* based on its size recovery. *S307* suppressed the small size phenotype of *sta1-1*, exhibiting the phenotype similar to wild-type (WT, Columbia harboring a *RD29A* promoter-driven luciferase transgene) (Figure 1A). In addition to the restored size, the splicing defects observed in *sta1-1* were also restored to wild-type levels in *S307* (Figures 1B and 1C). Intron retention (IR) profiles of *S307* were more similar to those of WT rather than to those of *sta1-1*, as shown by heatmaps (Figure 1B). Only 16 transcripts (16 increased, 0 decreased) displayed differential IR between *S307* and WT, in contrast to 2,876 transcripts (2,860 increased, 16 decreased) between s*ta1-1* and WT (Figure 1C, Supplemental Table 1).

**Figure 1.**
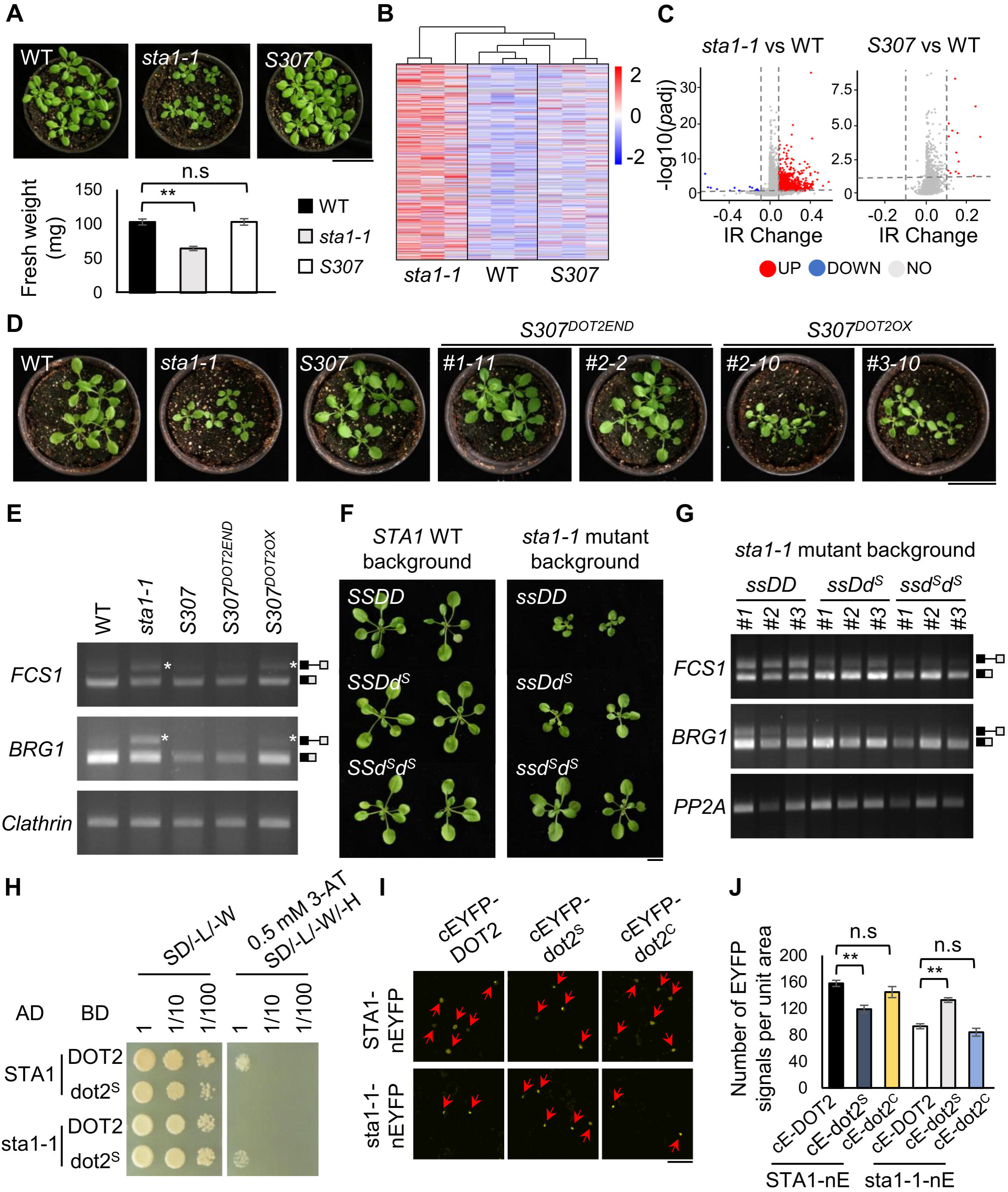
The *dot2^S^* mutation in *S307* rescues *sta1-1* defects and restores interaction with the sta1-1 protein. **(A)** Growth phenotypes and fresh weights of 3-week-old WT, *sta1-1*, and *S307* plants (scale bar = 5 cm; n = 22). **(B)** Heatmap showing IR ratios in WT, *sta1-1*, and *S307*. Three independent biological replicates were analyzed for each genotype. **(C)** Volcano plots showing significantly altered IR events. Blue and red dots represent significantly decreased and increased IR events, respectively. **(D)** Growth phenotypes of 3-week-old WT, *sta1-1*, *S307*, *S307^DOT2END^,* and *S307^DOT2OX^* plants (scale bar = 5 cm). **(E)** Experimental validation of IR levels for representative genes *FCS1* (*LEAFY CURLY AND SMALL1*, At1g78910), *BRG1* (*BOI-RELATED GENE1*, At5g45100), and *BGLU6* (*BETA GLUCOSIDASE6,* At1g60270). Asterisks indicate unspliced transcripts. Box diagrams show spliced and unspliced isoforms. *Clathrin* (At4g24550) is used as a loading control. **(F and G)** Semi-dominant effects of the *dot2^S^* mutation on plant growth and splicing in the *sta1-1* mutant background (scale bar = 1 cm). Three-week-old plants derived from an F2 population segregating for both the *sta1-1* and *dot2^S^* alleles (WT x *S307* cross). Genotypes are indicated as follows; *S*, *STA1*; *s, sta1-1*; *D, DOT2*; *d^S^*, *dot2^S^*. *PP2A* (*Protein phosphatase 2A*, At1g13320) is used as a loading control. Box diagrams showed spliced and unspliced isoforms. **(H)** Y2H assays demonstrating the interaction between STA1/sta1-1 and DOT2/dot2^S^. Yeast cells co-transformed with the fusion constructs of the GAL4 DNA-binding domain and activation domain were grown overnight and spotted onto selection medium (-L/-W), and interaction medium (-L/-W/-H) supplemented with 0.5 mM 3-AT (3-amino-1,2,4-triazole). **(I and J)** BiFC analysis for detecting the interaction between STA1/sta1-1 and DOT2/dot2^S^/dot2^C^ in leaves of *N*. *benthamiana* (scale bar = 75 μm). Red arrows indicate EYFP signals representing protein-protein interactions in the nucleus. EYFP signals were quantified from identical areas of leaf disks (2 x 2 cm). Data are presented as mean ± SE (n = 5). Statistical analysis is performed using one-way ANOVA with Tukey’s HSD test (\*\**P* < 0.01; n.s., not significant).

Whole-genome sequencing combined with map-based positional cloning using the F2 suppressor seedlings from a cross between *S307* and Landsberg *erecta* identified a G-to-A point mutation in the 8^th^ exon of the *DEFECTIVELY ORGANIZED TRIBUTARIES 2* (*DOT2*, At5g16780) gene. This mutation results in an amino acid change from aspartate to asparagine (Supplemental Figure 1). The *DOT2* gene encodes a protein homologous to Snu66p in *Saccharomyces cerevisiae* and SART1 in *Homo sapiens* (Supplemental Figure 2A), both of which are components of the U4/U6·U5 tri-snRNP together with STA1. Indeed, *DOT2* promoter-driven GUS activity was detected across all tissues, as found in the *STA1::GUS* plants (Lee et al., 2006) (Supplemental Figure 2B). This widespread and overlapping expression pattern suggests constant and cooperative roles for DOT2 and STA1.

To further examine whether DOT2 functions together with STA1, we characterized two independent *dot2* single mutant alleles: a *dot2* single mutant carrying the *S307* allele and a *dot2* T-DNA insertion mutant (SAIL_775_F10, also known as *mdf-2*) (de Luxan-Hernandez *et al*., 2022), hereafter referred to as *dot2^S^* and *dot2^T^*, respectively. *dot2^S^* plants were comparable to WT in both growth and splicing patterns, whereas *dot2^T^* plants showed growth and splicing defects resembling those of *sta1-1* (Supplemental Figures 3A and 3B). Consistent with previous reports (de Luxan-Hernandez *et al*., 2022), *dot2^T^*plants displayed cell death in the root apical meristem (RAM) as revealed by propidium iodide (PI)-stained confocal imaging. However, no detectable RAM defects were observed in the *dot2^S^* plants (Supplemental Figure 3C). Together, these results indicate that *dot2^S^* represents a weak loss-of-function allele and that *dot2* loss-of-function mutants display *sta1-1*-like defects, suggesting a coordinated function for DOT2 and STA1 in growth and pre-mRNA splicing.

### Genetic and molecular assays confirm *dot2^S^* as a semi-dominant suppressor of *sta1-1*

To confirm that the *dot2^S^* mutation is the causal mutation, a genomic fragment of *DOT2* including its endogenous promoter was introduced into *S307* via *Agrobacterium*-mediated transformation. Unexpectedly, transgenic plants expressing this genomic fragment of *DOT2* (*S307^DOT2END^*) did not revert the WT-like phenotype of *S307* to the small-sized *sta1-1* phenotypes (Figure 1D). However, when the *DOT2* gene was overexpressed by the Cauliflower Mosaic Virus (CaMV) 35S promoter in *S307* (*S307^DOT2OX^*), the overexpressing plants exhibited the *sta1-1*-like small size (Figure 1D). In addition, compared to *S307^DOT2END^, S307^DOT2OX^* exhibited higher accumulation of intron-retained transcripts for several marker genes that show IR in *sta1-1*, showing stronger phenotypic reversion by the overexpression (Figure 1E). We speculated that phenotypic reversion only by *DOT2* overexpression might be due to the dominant or semi-dominant effects of the dot2^S^ mutant protein on sta1-1, relative to those of the wild-type DOT2 protein; the dot2^S^ protein may interact more effectively with the sta1-1 protein than the DOT2 protein. Thus, we performed a dominance test on *dot2^S^* by genetically combining *dot2^S^* with the *STA1* homozygotes (WT) or the *sta1-1* homozygotes. No clear size defects were observed in the combinations of *dot2^S^*mutation in the wild-type background (Figure 1F). In contrast, the *dot2^S^* heterozygote in the *sta1-1* background showed an intermediate size, supporting the notion of semi-dominant effects of dot2^S^ on sta1-1 (Figure 1F). Consistently, the *dot2^S^* heterozygote in the *sta1-1* background exhibited intermediate levels of splicing defects between the *DOT2* (i.e., *sta1-1*) and the *dot2^S^* homozygotes in *sta1-1* (i.e., *S307*) (Figure 1G). Taken together, these results confirm that the mutation in *DOT2* is responsible for the *sta1-1* phenotypic suppression in *S307*.

### The *dot2^S^* mutation restores physical interaction with sta1-1, rescuing *sta1-1* defects

Homologs of STA1 and DOT2, as well as STA1 and DOT2 themselves, are known to interact, as supported by structural and experimental evidence (de Luxan-Hernandez *et al*., 2022; Häcker et al., 2008; Liu *et al*., 2006; Sun et al., 2018; Zhan *et al*., 2018). Our structural modeling predicted that the *sta1-1* and *dot2^S^* mutations are positioned near the STA1-DOT2 interaction interface (Supplemental Figure 4A). The *sta1-1* mutation resides within the HAT domain which mediates protein-protein interactions (Zeytuni and Zarivach, 2012), and the *dot2^S^* mutation maps to the predicted STA1-interacting surface identified by InterProSurf and PDBePISA (Krissinel and Henrick, 2007; Negi et al., 2007) (Supplemental Figure 4B). These findings suggest that both mutations may alter the STA1-DOT2 interaction. We examined interactions between the wild type and mutant proteins using yeast two-hybrid (Y2H) assays. As expected, STA1 and DOT2 showed substantial interaction (Figure 1H), whereas interactions between a wild-type protein (STA1 or DOT2) and a mutant protein (sta1-1 or dot2^S^) were not detectable. Interestingly, and consistent with our expectation, sta1-1 and dot2^S^ showed interactions comparable to those between STA1 and DOT2 (Figure 1H). Consistent with these observations, bimolecular fluorescence complementation (BiFC) analysis further confirmed the sta1-1-dot2^S^ interaction similar to the STA1-DOT2 interaction (Figures 1I and 1J).

To determine whether the suppressive effect is allele-specific, we generated an additional *dot2* allele using CRISPR/Cas9 (herein, called *dot2^C^*). The *dot2^C^*mutation is a 3-bp deletion located 44 bp upstream of the *dot2^S^* mutation, outside the predicted STA1-DOT2 interaction interface (Supplemental Figures 4A-4C). Unlike dot2^S^, the dot2^C^ protein did not alter interaction strength with STA1 or sta1-1 proteins (Figures 1I and 1J). Consistent with this, *dot2^C^* mutation did not alter the defects in size or intron retention observed in *sta1-1* (Supplemental Figures 4D and 4E). These results indicate that *dot2^S^* is a specific suppressor allele whose product interacts with the sta1-1 mutant protein, thereby likely restoring the U4/U6·U5 tri-snRNP assembly in *S307*.

### Interaction between STA1 and DOT2 promotes NS formation

We noticed diverse nuclear localization signals in our BiFC assays. Thus, we examined the localization of each GFP-tagged protein (STA1/sta1-1 and DOT2/dot2^S^/dot2^C^) and classified the nuclear signals into five subclasses based on their predominant subnuclear compartments (Figure 2A). Our observation revealed that STA1-GFP formed nuclear condensates (Class IV and V) at a higher frequency than sta1-1-GFP, whereas GFP-DOT2/dot2^S^/dot2^C^ were predominantly localized to the nucleolus (Class II) (Figure 2B). Interestingly, co-expression of STA1-GFP and mRFP-DOT2 significantly increased condensate formation of both proteins compared to expression of each protein alone (from ∼15%/0% to ∼92% for STA1/DOT2) (Figures 2A-2E). This suggests that the STA1-DOT2 interaction contributes to the localization of each protein to nuclear condensates. This idea was supported by the following observations: sta1-1-GFP showed limited condensate formation when co-expressed with mRFP-DOT2/dot2^C^, whereas mRFP-dot2^S^ strongly enhanced sta1-1-GFP condensate formation (Figures 2C-2E), consistent with the interaction strengths of these protein combinations. To identify the nature of the nuclear condensates observed above, we co-expressed each protein with the NS marker mRFP-U1-70K (Ali et al., 2008). STA1-GFP condensates formed upon single expression of STA1-GFP showed partial co-localization with mRFP-U1-70K (Figure 2F). Upon co-expression with DOT2, STA1-GFP became nearly completely co-localized with mRFP-U1-70K (Figure 2G). Furthermore, the EYFP signals from the BiFC assay (i.e., co-expression of STA1 and DOT2), indicating the STA1-DOT2 interaction, showed complete co-localization with mRFP-U1-70K (Figure 2H). These results confirm that the STA1-DOT2 interaction promotes the formation of nuclear condensates corresponding to NS. NS are a specific type of nuclear condensates that are enriched in the spliceosome components and associated with mRNA splicing and mRNA metabolism (Bhat et al., 2024; Daguenet *et al*., 2012). Therefore, our results indicate that the STA1-DOT2 interaction contributes to their NS localization and probably to proper mRNA splicing. It should be noted that STA1-GFP condensates that did not overlap with mRFP-U1-70K suggest that STA1-GFP may form non-functional or NS-like condensates (herein called “aggregates”) (see below).

**Figure 2.**
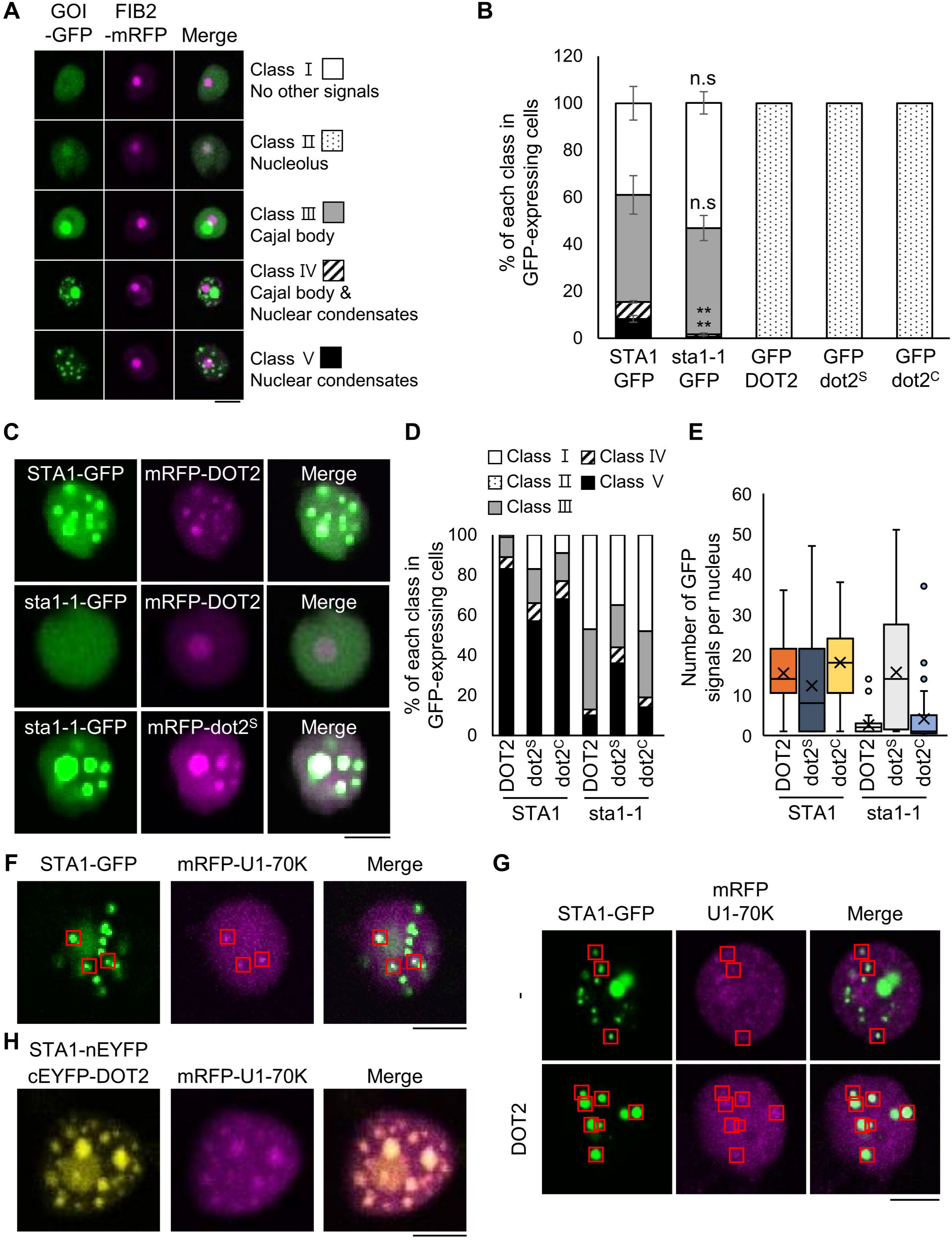
The disrupted nuclear localization of sta1-1 protein is re-localized to NS by the dot2^S^ protein. **(A)** Classification of nuclear compartments showing the localization of GOI (gene of interest)-GFP driven by the CaMV 35S promoter into five subclasses. The nucleolus marker FIB2 (FIBRILLARIN 2, At4g25630)-mRFP was co-infiltrated (scale bar = 10 μm). **(B)** Subcellular localization of each protein tagged with GFP in leaves of *N. benthamiana*. Nuclear localization patterns of each GFP-tagged protein were classified into the five distinct categories defined in **(A)**. For each set, 100 GFP-expressing cells were analyzed per replicate, and data are presented as mean ± SE (n = 4). Statistical analysis is performed using one-way ANOVA with Tukey’s HSD test (\*\**P* < 0.01; n.s., not significant). **(C-E)** Co-localization of GFP-tagged STA1/sta1-1 with mRFP-tagged DOT2/dot2^S^/dot2^C^ in leaves of *N. benthamiana*. Nuclear localization categories are shown in **(D)**, classified according to the groups defined in **(A)**. The number of NS per nucleus is shown in **(E)**, based on the STA1/sta1-1-GFP signals upon co-expression with mRFP-DOT2/dot2^S^/dot2^C^. For each analysis, 100 cells **(D)** or 50 nuclei **(E)** were randomly selected and analyzed (scale bar = 10 μm). **(F-H)** Localization of GFP-tagged STA1 or BiFC signals of STA1 and DOT2 in leaves of *N. benthamiana*. Co-localization of GFP-tagged STA1 **(F and G),** upon single or co-expression with DOT2, and EYFP signals from BiFC analysis of the STA1 and DOT2 interaction **(H)** with the NS marker U1-70K (U1 SMALL NUCLEAR RIBONUCLEOPROTEIN-70K, At3g50670) tagged with mRFP (mRFP-U1-70K). Red boxes indicate regions of overlap between STA1-GFP/BiFC signals and mRFP-U1-70K (scale bar = 10 μm). No red boxes are indicated in **(H)** as the signals overlap completely.

### The *dot2^S^* mutation restores sta1-1 localization to NS, enabling proper splicing

We next examined the functional significance of NS formation of STA1 and DOT2 for mRNA splicing. To this end, we generated the STA1-GFP/sta1-1-GFP overexpressing plants in *Salk007933*, a *sta1* T-DNA insertion knock-out line (hereafter referred to as *Salk007933^STA1-GFP^*and *Salk007933^sta1-1-GFP^*, respectively). Homozygous *sta1* knockouts are lethal. However, introduction of either STA1-GFP or sta1-1-GFP rescued this lethality and the resulting plants phenocopied the wild-type and *sta1-1* plants, respectively (Supplemental Figure 5). Using protoplasts isolated from these GFP-expressing plants, we confirmed again that the interaction between STA1 and DOT2 (or sta1-1 and dot2^S^) promoted the localization of the GFP-tagged protein to NS (Figure 3A).

**Figure 3.**
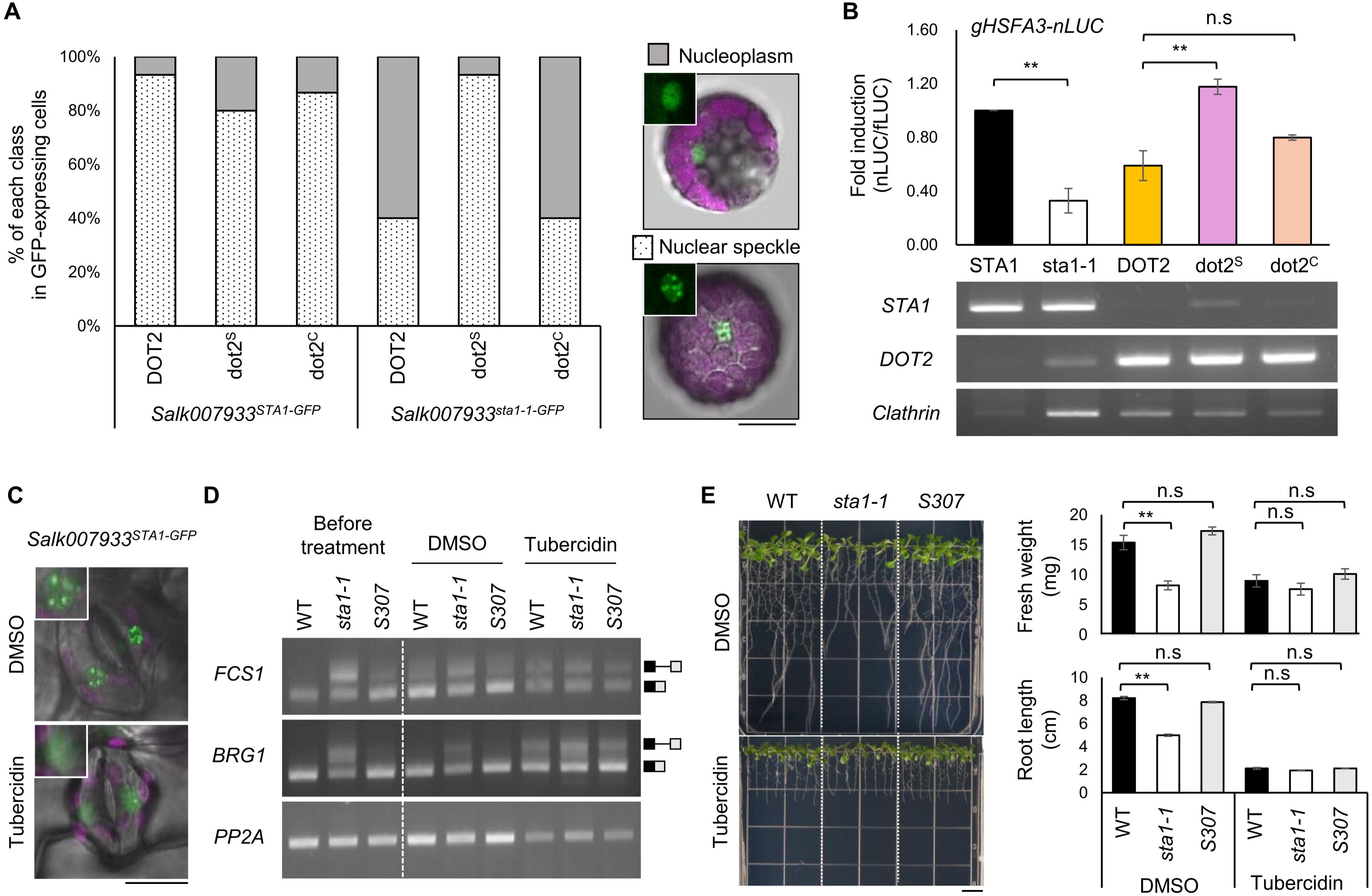
The localization of STA1 to NS is closely associated with proper pre-mRNA splicing. **(A)** Nuclear localization patterns of GFP-tagged *STA1*/*sta1-1* upon transfection with *DOT2*/*dot2^S^*/*dot2^C^*in *Salk007933^STA1-GFP^* or *Salk007933^sta1-1-GFP^*protoplasts. Cells with nuclear-localized GFP signals of two distinct patterns (nucleoplasm and NS) were quantified as a ratio. A total of 15 cells were analyzed for each combination (scale bar = 5 μm). **(B)** Splicing activity of *DOT2*/*dot2^S^*/*dot2^C^*on the *gHSFA3* DNA in *Salk007933^sta1-1-GFP^* protoplasts. Splicing activity on the *gHSFA3* DNA is represented as the relative luminescence intensity by NanoLuc luciferase (nLUC) normalized to firefly luciferase (fLUC). *Clathrin* is used as a loading control. Data are presented as mean ± SE (n = 2). Statistical analysis is performed using one-way ANOVA with Tukey’s HSD test (\*\**P* < 0.01; n.s., not significant). **(C)** Tubercidin-induced delocalization of STA1 from NS to nucleoplasm. Ten-day-old *Salk007933^STA1-GFP^* plants were treated with DMSO or 100 μM tubercidin for 1 h (scale bar = 25 μm). **(D)** Tubercidin-induced disruption of pre-mRNA splicing. Ten-day-old WT, *sta1-1*, and *S307* plants were treated with DMSO or 100 μM tubercidin for 12 h. The box diagrams show spliced and unspliced isoforms. *PP2A* is used as a loading control. **(E)** Abolished size differences between WT, *sta1-1*, and *S307* by tubercidin treatment (scale bar = 1 cm). Five-day-old seedlings were transferred to medium containing either DMSO or 10 μM tubercidin for 6 d. Data are presented as mean ± SE (n = 4).

Previously, we developed a protoplast-based splicing activity assay using a luciferase reporter gene fused with a genomic fragment of *HSFA3* (*gHSFA3-LUC*) which undergoes STA1-dependent splicing (Kim *et al*., 2017). After transfection with the alleles to be tested, splicing activity was measured in protoplasts derived from *Salk007933^sta1-1-GFP^*. As expected, the introduction of *STA1* into the protoplasts successfully restored full splicing activity of *gHSFA3-LUC*, while *sta1-1* did not (Figure 3B). Among the tested *DOT2* alleles, only the *dot2^S^*allele showed the full splicing activity in *Salk007933^sta1-1-GFP^* (Figure 3B). Taken together, these results indicate that the STA1-DOT2 or sta1-1-dot2^S^ interaction is critical for NS formation and splicing activity, thereby linking NS formation to proper splicing activity.

We then examined the effect of disrupting NS formation by applying tubercidin, an inhibitor of speckle formation (Kurogi et al., 2014) to *Salk007933^STA1-GFP^* plants. We observed that STA1-GFP in *Salk007933^STA1-GFP^* was localized predominantly to NS (Figure 3C). However, tubercidin treatment dispersed the GFP signals into the nucleoplasm (Figure 3C). Tubercidin treatment also eliminated the differences in splicing activity and growth patterns among WT, *sta1-1*, and *S307*, resulting in comparable splicing and growth phenotypes across all genotypes (Figures 3D and 3E). Altogether, these results demonstrate that the interaction-dependent localization of STA1 and DOT2 is functionally critical for proper mRNA splicing and that the *S307* suppressor phenotypes are due to the rescued interaction between sta1-1 and dot2^S^.

### Heat-induced weakening of the STA1–DOT2 interaction reduces nuclear speckle formation

To further assess the functional connection between the STA1-DOT2 interaction and speckle formation, we employed heat as an additional perturbation to weaken protein-protein interactions, because elevated temperatures can weaken molecular interactions. We also hypothesized that the heat-hypersensitive phenotype of *sta1-1* might result from heat-induced weakening of the sta1-1 interaction. Consistent with this idea, BiFC analysis revealed a reduction in EYFP signals corresponding to the STA1-DOT2 interaction after heat treatment (Figures 4A-4C).

**Figure 4.**
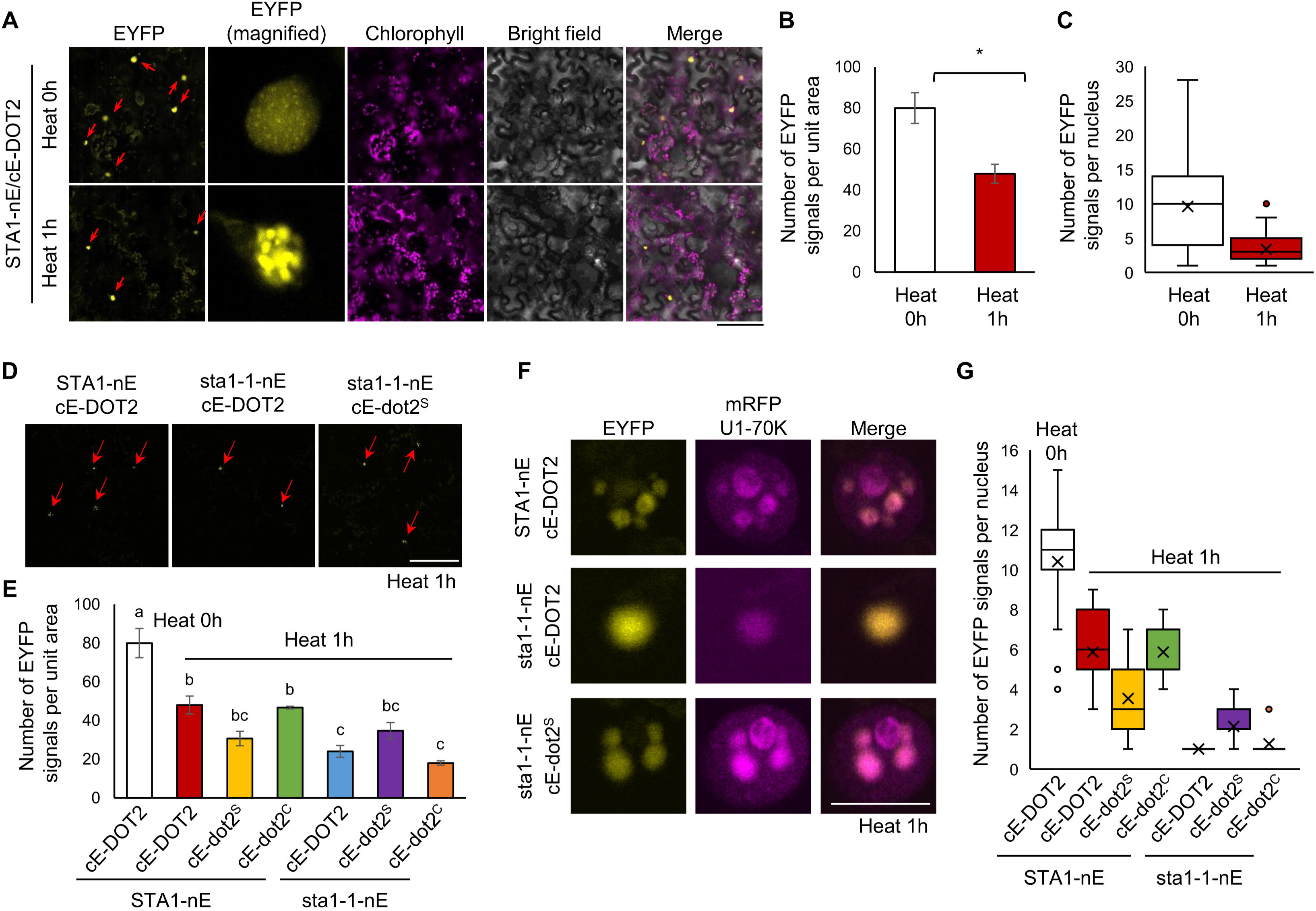
The dot2^S^ protein facilitates NS localization of the sta1-1 protein under heat conditions. **(A-C)** BiFC analysis to detect interaction between STA1 and DOT2 under heat conditions in leaves of *N. benthamiana* (scale bar = 75 μm). Red arrows indicate EYFP signals representing STA1-DOT2 interactions in the nucleus. The number of EYFP signals per unit area (2 x 2 cm) and per nucleus (n = 43) was quantified in **(B)** and **(C)**, respectively. **(D and E)** BiFC analysis to detect interaction between STA1/sta1-1 and DOT2/dot2^S^/dot2^C^ under heat stress conditions in leaves of *N. benthamiana* (scale bar = 75 μm). Red arrows indicate EYFP signals. EYFP signals were quantified from the same area of leaf disks (1 x 1 cm). Data are presented as means from three independent biological replicates. Statistical analysis is performed using one-way ANOVA with Tukey’s HSD test. Significant differences are denoted by letters above the plots. **(F and G)** Co-localization of EYFP signals from BiFC analysis of STA1/sta1-1 and DOT2/dot2^S^/dot2^C^ with the NS marker mRFP-U1-70K under heat stress conditions in leaves of *N. benthamiana* (scale bar = 10 μm). Box plots show the number of NS exhibiting EYFP signals. Center lines indicate medians; boxes represent interquartile range, and whiskers extend to minimum and maximum values. For heat treatment, samples were incubated at 37°C for 1 h in liquid MS medium following a 36 h post-infiltration incubation.

Unexpectedly, despite the reduction in BiFC signals, heat treatment increased the numbers of STA1-GFP condensates when STA1-GFP was expressed alone (Supplemental Figure 6A). We hypothesized that the increase of heat-induced condensates might result from STA1 self-interaction (i.e., STA1 aggregation). To test this possibility, we performed BiFC analysis using STA1-nEYFP and STA1-cEYFP. Heat treatment indeed increased BiFC signals, indicating enhanced STA1 self-interaction under heat (Supplemental Figure 6B). Most EYFP signals derived from STA1 self-interaction did not overlap with signals from the speckle marker mRFP-U1-70K under either normal or heat conditions (Supplemental Figures 6C and 6D), suggesting that the STA1 self-interacting condensates represent STA1 aggregates. This interpretation is further supported by the presence of predicted intrinsically disordered regions (IDRs) within STA1 (residues 109-216, 298-318, 910-918, and 1022-1029), as identified by MolPhase (Liang et al., 2024) (Supplemental Figure 6E). Therefore, the increase in STA1-GFP condensates observed under STA1-GFP single expression is likely the result of STA1-GFP self-aggregation.

To distinguish the NS from STA1 aggregates, we used BiFC analysis to quantify NS induced by the STA1-DOT2 interaction. The number of these speckle-representing BiFC signals in nuclei decreased after heat treatment and speckle formation was constantly lower in combinations of the weakly interacting STA1 and DOT2 allele products (Figures 4D-4G). These results reaffirm that the STA1-DOT2 interaction is required for efficient NS formation.

### The *dot2^S^* mutation suppresses the heat sensitivity of *sta1-1*

Next, we sought to determine whether the strength of the STA1-DOT2 interaction also plays an important role in the heat responses. The *sta1-1* mutants have previously been shown to display heat-sensitive phenotypes (Kim *et al*., 2017; Kim et al., 2018). At both the seedling and reproductive stages, the heat-sensitive phenotypes of *sta1-1* were almost fully rescued in *S307* (Figures 5A–5C). Together, these results, along with the partial or full reversion to *sta1-1* heat sensitivity observed in *S307^DOT2OX^*, indicate that DOT2 functions together with STA1 under heat stress (Figures 5A and 5C). In addition, the *dot2^C^* allele did not suppress the heat sensitivity of s*ta1-1*, supporting the allele-specific suppressive effect of *dot2^S^*(Figures 5B and 5C). These findings also suggest that the restored heat tolerance in *S307* results from the rescued interaction between sta1-1 and dot2^S^.

**Figure 5.**
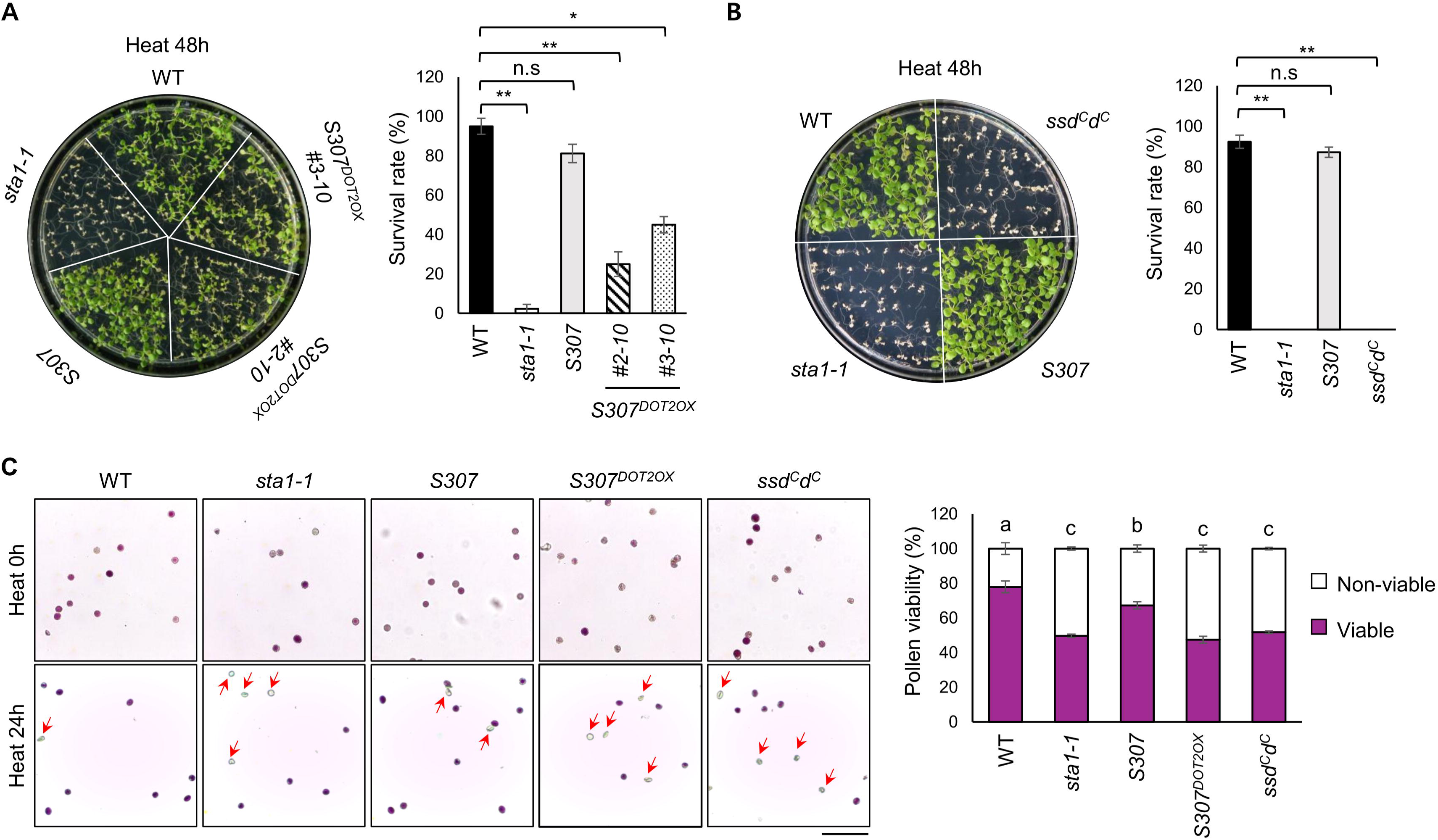
Heat-sensitivity of *sta1-1* is restored to wild type levels in the *S307* suppressor. **(A and B)** Heat survival test and quantification of survival rate of each plant genotype. Ten-day-old plants of each genotype were exposed to heat stress (37°C for 48 h), and survived plants were scored. Statistical analysis is performed using one-way ANOVA with Tukey’s HSD test (\*\**P* < 0.01; \**P* < 0.05; n.s., not significant). **(C)** Assessment of pollen viability using Alexander staining (scale bar = 50 μm). Viable pollen grains appear purple, while non-viable pollen grains are transparent. Red arrows indicate non-viable pollen grains. Data are presented as means ± SE from three independent biological replicates. Statistical analysis is performed using one-way ANOVA with Tukey’s HSD test. Significant differences are denoted by letters above the plots.

### The *dot2^S^* mutation mitigates transcriptome-wide intron retention defects in *sta1-1* under heat stress

Under heat stress, IR profiles in *S307* were partially restored toward WT levels (Figure 6A). Although the global splicing patterns in *S307* did not fully return to wild-type levels, the IR defects of *sta1-1* were markedly improved in *S307*. Among 67,103 transcripts with significant IR changes across genotypes, *S307* showed a reduced median IR ratio (0.14) compared with *sta1-1* (0.34) (Figure 6B). Across the three genotypes under heat, 1,114 introns (from 940 genes) showed increased IR in WT, whereas 58,522 introns (12,140 genes) and 36,617 introns (10,654 genes) showed increased IR in *sta1-1* and *S307*, respectively (Figure 6C, Supplemental Table 2). Venn diagram analysis further revealed that 27,514 of the 58,522 introns (∼47%) with elevated IR in *sta1-1* did not exhibit increased IR in *S307* (Figure 6D). Together, these results indicate a partial restoration of *sta1-1* splicing defects in *S307* under heat stress.

**Figure 6.**
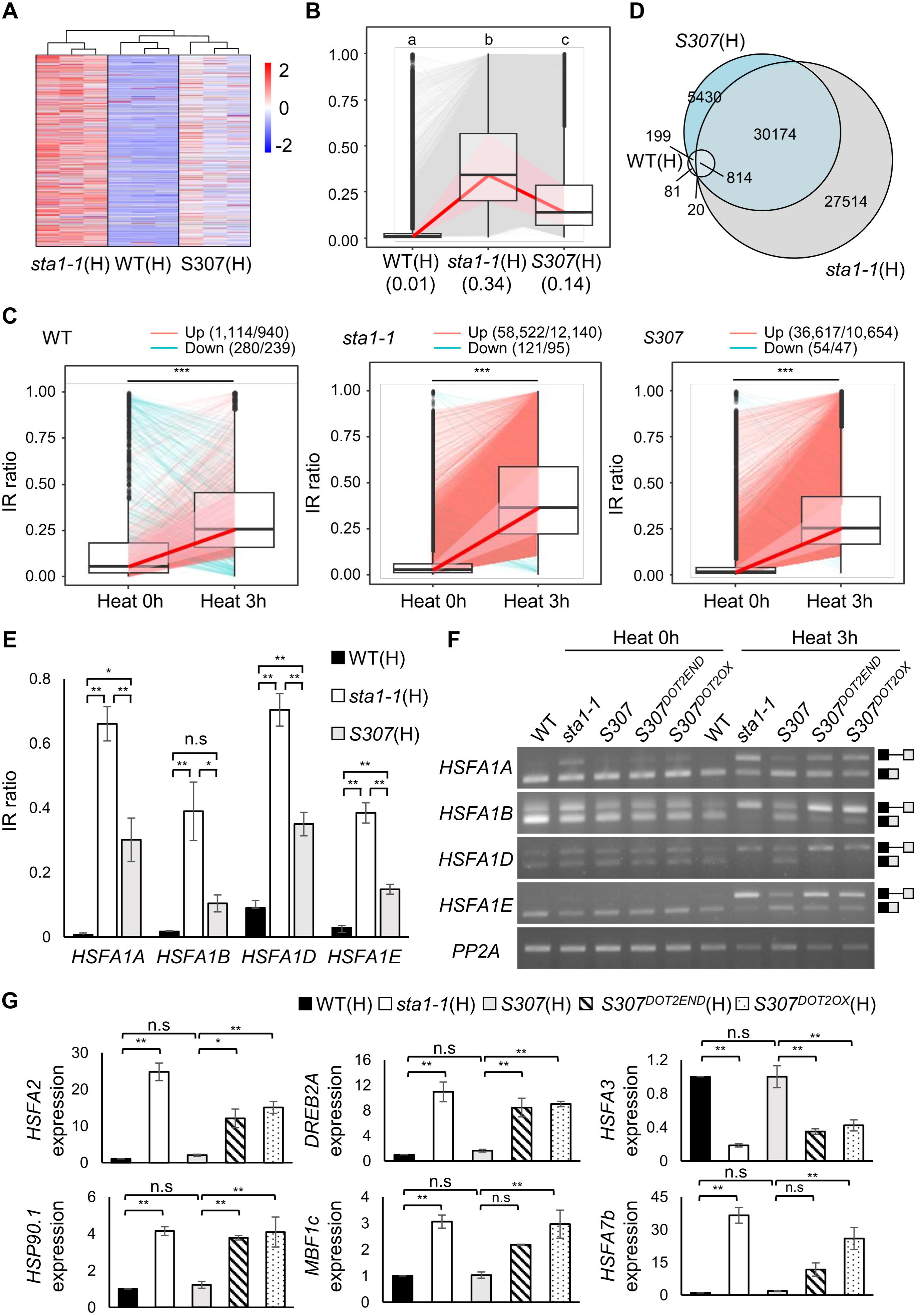
Defective splicing of *HSFA1s* in *sta1-1* is rescued in *S307*, leading to proper expression of downstream target genes. **(A)** Heatmap showing the IR ratios in WT under heat stress (WT(H)), *sta1-1* under heat stress (*sta1-1*(H)), and *S307* under heat stress (*S307*(H)). Each genotype is represented by three independent biological replicates. **(B)** Boxplot analysis showing IR ratio trends across genotypes for 67,103 transcripts exhibiting significant changes in intron retention. Each box represents the IR ratio distribution for each genotype, with lines connecting genotypes to indicate intron-specific changes. The ribbon highlights the median and variability for visual summary, and the bold red line indicates the overall trend of median IR ratios across genotypes. Median IR ratios for each genotype are shown in parentheses. Statistical significance among genotypes was assessed using the Kruskal–Wallis test followed by a post hoc Dunn’s test with Benjamini–Hochberg correction. Significant differences are denoted by letters above the plots. **(C)** Changes in intron retention levels of transcripts in response to heat stress across different genotypes. Paired notation denotes (number of introns/number of genes). **(D)** Venn diagram showing transcripts with increased intron retention under heat stress in each genotype. **(E and F)** Quantification and validation of splicing defects in the master heat stress regulators *HSFA1*s. Box diagrams show spliced and unspliced isoforms. *PP2A* is used as a loading control. **(G)** qRT-PCR analysis of heat stress-related genes regulated by *HSFA1*s. The relative expression levels of each gene are shown in the graphs. Data are presented as mean ± SE from three independent biological replicates. Statistical analysis is performed using one-way ANOVA with Tukey’s HSD test (\*\**P* < 0.01; \**P* < 0.05; n.s., not significant).

Among the genes with restored splicing patterns, *HSFA1*s (*HEAT SHOCK TRANSCRIPTION FACTOR A1*s), key upstream regulators of heat-responsive genes (Li et al., 2023; Nishizawa-Yokoi et al., 2011; Röth et al., 2017), showed reduced intron retention in *S307* compared with *sta1-1* (Figure 6E, Supplemental Figure 7). This recovery was further supported by decreased levels of spliced transcripts and increased levels of intron-retained transcripts of *HSFA1*s in *S307^DOT2END^* and *S307^DOT2OX^* relative to *S307* (Figure 6F). The *dot2^C^*allele failed to rescue splicing defects of *HSFA1s* in *sta1-1*, consistent with its heat-sensitive phenotypes (Figures 5B and 5C, Supplemental Figure 8). Proper splicing of *HSFA1s* in *S307* enabled normal expression of downstream heat-responsive genes (Figure 6G). These findings indicate that *dot2^S^* suppresses intron retention defects in heat-responsive genes in *sta1-1*, thereby contributing to the restoration of heat tolerance. Taken together with the previous results, these findings highlight the importance of the STA1-DOT2 interaction for proper heat stress responses through NS formation and accurate mRNA splicing under heat conditions. The incomplete rescue of splicing defects and heat sensitivity in *S307* under heat stress, particularly at the pollen stage (Figure 5C), is likely attributable to the weaker sta1-1-dot2^S^ interaction compared with the wild-type STA1-DOT2 interaction.

## DISCUSSION

Nuclear speckles (NS) are dynamic, membraneless nuclear compartments enriched in spliceosomal components and RNAs (Mintz et al., 1999; Smith *et al*., 2020), and are increasingly recognized as active hubs that facilitate efficient pre-mRNA processing rather than acting merely as passive storage sites (Wu *et al*., 2024). Previous studies have emphasized the contribution of intrinsically disordered regions, low-complexity domains, and multivalent RNA–protein interactions to NS organization (Ku et al., 2025; Paul *et al*., 2024; Wang *et al*., 2024; Zhang et al., 2024). To what extent spliceosomal protein-protein interactions involved in spliceosome assembly contribute to NS formation remains poorly understood. In this study, we provide evidence that NS organization in plants is tightly associated with the integrity of a specific interaction between two core U4/U6·U5 tri-snRNP components, STA1 and DOT2, thereby linking spliceosome assembly with subnuclear organelle organization.

Our genetic and cell biological analyses show that the weakening of the STA1-DOT2 interaction by the *sta1-1* mutation is accompanied by reduced NS formation, compromised splicing efficiency, and growth defects, whereas restoration of this interaction by the *dot2^S^* suppressor allele is associated with recovery of these phenotypes. Importantly, this suppression is achieved through reinforcement of the STA1-DOT2 interaction probably within the tri-snRNP. These observations suggest that NS organization is not independent of spliceosome assembly status and is closely linked to the stability of specific spliceosomal interactions.

The spliceosome is a highly dynamic macromolecular complex, and formation of the U4/U6·U5 tri-snRNP represents a critical step in each splicing cycle (Will and Lührmann, 2011). PRP6/STA1 functions as a central bridging factor between the U4/U6 di-snRNP and the U5 snRNP, with its interactions stabilized by tri-snRNP–specific proteins such as SART1/DOT2 (Bertram *et al*., 2017; Nguyen *et al*., 2016). Our observation of a positive correlation between STA1–DOT2 interaction strength and NS abundance supports a model in which proper tri-snRNP assembly facilitates efficient recruitment or retention of mature spliceosomal complexes within NS. Consistent with this view, quantitative subnuclear localization analyses revealed that co-expression of STA1-GFP and mRFP-DOT2 significantly increased NS formation (types IV and V), compared with expression of each protein alone (Figures 2B and 2D). In contrast, co-expression of sta1-1-GFP and mRFP-DOT2 resulted in reduced NS-associated localization patterns (types IV and V) and increased localization to Cajal bodies (type III) compared with the wild-type protein (Figure 2D). Tri-snRNPs are assembled in Cajal bodies and subsequently redistributed to NS, which contain high levels of mature, splicing-competent complexes (Schaffert *et al*., 2004; Yildirim *et al*., 2021). Thus, our results suggest that the *sta1-1* mutation impairs later steps associated with NS organization due to defects in the STA1–DOT2 interaction, likely leading to impaired formation of splicing-competent spliceosome. In agreement with our results and speculation, when SART1, homologous to DOT2, was immunologically depleted from HeLa cells, pre-mRNA splicing was inhibited but the integrity of the U4/U6·U5 tri-snRNP was not affected, indicating the importance of SART1 in active spliceosome formation and recruitment of the tri-snRNP to the pre-spliceosome (Makarova et al., 2001; Utans et al., 1992). Thus, the STA1–DOT2 interaction is likely to promote DOT2-dependent localization of STA1 to NS, concomitant with the delivery of the tri-snRNP to the pre-spliceosome for active spliceosome assembly.

The functional relevance of our notion becomes particularly apparent under heat stress conditions. Elevated temperatures are known to destabilize spliceosomal assemblies and broadly impair splicing fidelity (Bond, 1988; Bond and James, 2000; Park et al., 2024). The most sensitive component of the spliceosome is the U4/U6·U5 tri-snRNP (Bond, 1988). In this context, the STA1–DOT2 interaction appears to function as a heat-sensitive interaction node, since the heat-induced weakening of this interaction correlates with reduced NS formation, increased intron retention, and weakened heat tolerance (Figure 4, Figure 5, Figure 6). Reinforcement of the interaction by the dot2^S^ allele in sta1-1 (i.e., the *S307* suppressor) is associated with improved NS formation and splicing performance, as well as partial recovery of survival under heat stress (Figure 4, Figure 5, Figure 6). Although it is still debated whether NS primarily function as storage sites or as active hubs for RNA metabolism, our findings support the latter view by linking speckle organization to both defined molecular interactions and functional splicing outcomes. It should also be noted that STA1 itself appeared to form non-functional aggregates under heat conditions due to its IDRs (Supplemental Figure 6). DOT2 seemed to inhibit the aggregation of STA1 and direct STA1 to NS, presumably facilitating formation of an active spliceosome. Thus, STA1–DOT2 interaction contributes to the robustness of pre-mRNA splicing under heat stress.

Taken together, our study expands current understanding of NS biology by demonstrating that NS organization in plants is closely associated with the assembly state of the spliceosome, mediated by specific protein–protein interactions within the U4/U6·U5 tri-snRNP. On this basis, we propose that heat-sensitive interaction nodes such as the STA1–DOT2 bridge represent “molecular weak points” or “interaction-level sensors” that integrate spliceosome architecture, subnuclear organelle formation, and environmental responsiveness (Figure 7). This interaction-based perspective provides insight into how plants maintain splicing robustness under stress conditions and suggests that targeted modulation of spliceosomal interactions may offer a potential strategy for improving stress tolerance.

**Figure 7.**
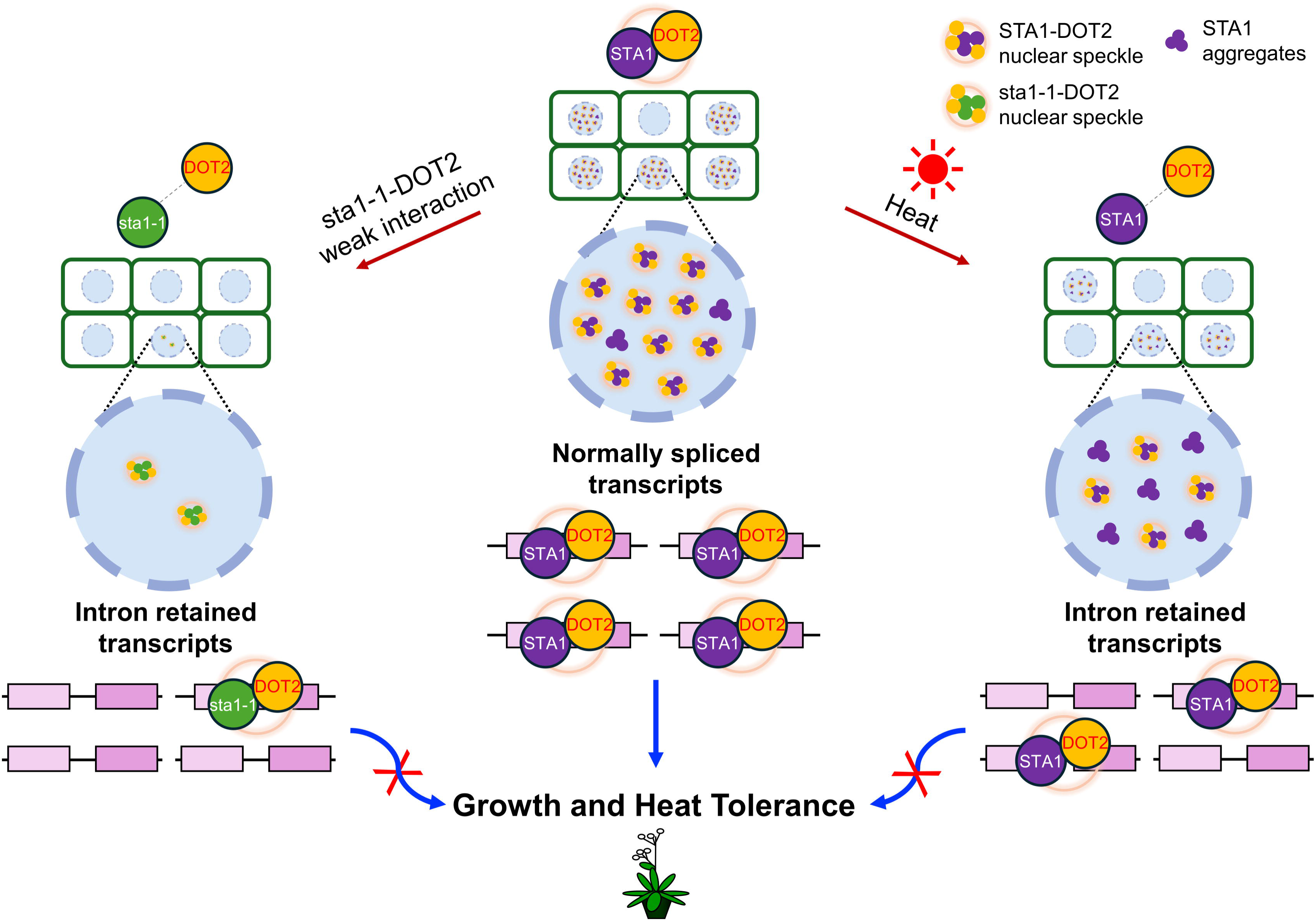
A proposed model for STA1-DOT2 interaction-mediated NS formation and pre-mRNA splicing in growth and heat stress responses. The STA1-DOT2 interaction, presumably reflecting the U4/U6·U5 tri-snRNP assembly, is important for NS formation. Proper NS formation ensures the production of correctly spliced transcripts required for normal growth and heat stress responses. The *sta1-1* mutation or heat stress weakens the STA1-DOT2 interaction, thereby reducing NS formation. This disrupted subnuclear localization of splicing components increases intron-retained transcripts, resulting in defective growth and heat stress responses.

## METHODS

### Plant growth conditions

*Arabidopsis thaliana* seeds were planted on Murashige and Skoog (MS) medium containing MS salts with vitamins (Caisson Laboratories Inc, USA), 2% sucrose, and 0.3% gelite (Duchefa, the Netherlands), adjusted to pH 5.7–5.8. For vertical growth, 0.6% gelrite was supplemented. Seeds were surface-sterilized with commercial bleach supplemented with 0.01% Tween 20 (USB, USA), stratified at 4°C for 2 days in the dark, and then transferred to a growth chamber under long-day conditions (16 h light/8 h dark) at 22°C. For soil-grown plants, seeds were sown on Sun Gro propagation mix (Sun Gro, Canada) and maintained under long-day conditions (16 h light/8 h dark) at 22°C.

### Positional cloning and mutation identification

To map the *S307* mutation locus, *S307* (Columbia background) was crossed with Landsberg *erecta* (L*er*) to generate an F2 mapping population. WT-sized F2 plants homozygous for the *sta1-1* mutation were selected, and recovery of the size phenotype was confirmed in the F3 population pool. Genomic DNA was extracted from pooled leaves of 94 selected F2 mutant individuals using the RNeasy Plant Mini Kit (QIAGEN, Germany), and whole-genome sequencing was performed at Macrogen Inc. The genomic region between the SSLP markers Ch5_5.46 Mb and Ch5_5.71 Mb, identified by initial mapping, was compared with the WT genomic sequence, identifying a candidate point mutation at position 1,651 bp of *DOT2* (At5g16780) coding sequence. The candidate mutation was validated by complementation using genomic constructs driven by either the native *DOT2* promoter (DOT2::DOT2) or the Cauliflower mosaic virus 35S promoter (35S::DOT2). For native *DOT2* promoter complementation lines (*S307^DOT2END^*), a 3,783 bp genomic fragment spanning 1,695 bp upstream of the *DOT2* start codon to 294 bp downstream of the stop codon was amplified and cloned into pCAMBIA1300 between the *Kpn*I and *Bam*HI sites (Takara, Japan). For 35S-driven complementation lines (*S307^DOT2OX^*), the 2,463 bp *DOT2* coding sequence was amplified, cloned into pDONR221 entry vector, and subsequently recombined into the pMDC32 destination vector using the Gateway cloning system. All primers used to generate complementation constructs are listed in Supplemental Table 3.

### Generation of different alleles of *dot2* mutants

The *dot2* single mutant carrying the *S307* allele (*dot2^S^*) was obtained from the F2 progeny of a cross between WT and *S307*. A derived cleaved amplified polymorphic sequence (dCAPS) marker detecting the single-nucleotide substitution in the *dot2^S^* allele using *Eco*RI (Takara, Japan), together with a marker detecting the 6-nucleotide deletion in *sta1-1* (used to exclude plants carrying the *sta1-1* mutation), was employed to isolate the *dot2^S^* single mutant. The *dot2* T-DNA insertion mutant *SAIL_775_F10* (*dot2^T^*) was obtained from the Arabidopsis Biological Resource Center (ABRC). Because homozygous *dot2^T^*plants are infertile, only heterozygotes were maintained, and homozygotes were identified by their dwarf phenotype (Petricka *et al*., 2008).

A CRISPR/Cas9-generated *dot2* mutant allele (*dot2^C^*) was produced in the *sta1-1* background using the CRISPR/Cas9 system as previously described (Cho et al., 2017; Liu et al., 2015; Yun et al., 2022). A single guide RNA (sgRNA) targeting *DOT2* was designed using the CRISPR web toolbox CHOPCHOP (https://chopchop.cbu.uib.no) (Labun et al., 2019). The annealed oligonucleotides (DOT2sgRNA3_infu_CF and DOT2sgRNA3_infu_CR) for sgRNA were cloned into the psgR-Cas9-At to generate *DOT2*-targeting construct. The resultant DOT2sgRNA3-Cas9 cassette, excised with *Hin*dIII and *Eco*RI (Takara, Japan), was subsequently subcloned into the *Hin*dIII-*Eco*RI sites of pCAMBIA1300. *Agrobacterium tumefaciens* strain GV3101 carrying these constructs was used to transform *sta1-1* plants by the floral dip method to obtain T1 seeds. CRISPR/Cas9-edited elite lines were identified by Sanger sequencing of the DOT2sgRNA3-targeted region in the T1 generation, and homozygous lines were subsequently isolated. All primers used for the isolation of different *dot2* mutant allels are listed in Supplemental Table 3.

### RNA extraction and cDNA synthesis

Total RNA was extracted from seedlings at the designated developmental stage using RNAiso Plus reagent (Takara, Japan) and treated with RNase-free DNase (New England BioLabs, USA). cDNA was synthesized from total RNA using the TOPscript^TM^ Reverse Transcriptase kit (Enzynomics, Korea). Semi-quantitative RT-PCR and realtime PCR analyses were performed using 2 μL of the resulting cDNA (corresponding to 50 ng of input RNA). All primers used for intron retention analyses and gene expression quantification are listed in Supplemental Table 3.

### GUS staining assay

For the *STA1* promoter-driven *GUS* (*STA1::GUS*) and *DOT2* promoter-driven *GUS* (*DOT2::GUS*) constructs, 1,475 bp and 1,695 bp promoter fragments upstream of the start codon of each gene were amplified, respectively. These fragments were cloned into pCAMBIA1381 between the *Bam*HI and *Hind*III sites for *STA1::GUS*, or between the *Bam*HI and *Nco*I sites for *DOT2::GUS* (Takara, Japan). *Agrobacterium tumefaciens* strain GV3101 carrying these constructs was used to transform Col-0 plants by the floral dip method.

Tissues of transgenic *Arabidopsis* seedlings carrying the *STA1::GUS* or *DOT2::GUS* constructs were collected in 90% acetone (Duksan, Korea) in conical tubes on ice and incubated for 10–20 min at room temperature. After removal of acetone, samples were washed twice for 5 min with washing buffer (50 mM potassium phosphate buffer (pH = 7.0), 0.2% Triton X-100, 2 mM potassium ferricyanide, and 2 mM potassium ferrocyanide). The washing buffer was then replaced with staining buffer (washing buffer supplemented with 2 mM X-Gluc (Sigma, Japan)), and tissues were incubated overnight at 37°C. Subsequently, tissues were incubated in 70% ethanol to remove chlorophyll. GUS-staining was observed using a light microscope (Leica ICC50, Germany).

### Nuclear localization, co-localization, and BiFC analyses

For nuclear localization analyses, the full-length coding regions of *STA1* (*sta1-1*) lacking a stop codon and *DOT2* (*dot2^S^*and *dot2^C^*) containing a stop codon were amplified, cloned into pDONR221 entry vector, and subsequently recombined into the pGWB551 or pGWB552 destination vectors, respectively, using the Gateway cloning system. *Agrobacterium tumefaciens* cells harboring *STA1 (sta1-1)-GFP* or *GFP-DOT2* (*dot2^S^* and *dot2^C^*) constructs were infiltrated into *Nicotiana benthamiana* leaves in expression buffer (50 mM MES (pH = 5.7), 10 mM MgSO_4_, 5% D-Glucose, and 0.1 mM of acetosyringone) together with P19, each at a final concentration of OD_600_ of 0.2. After 36 h of incubation of infiltrated *N. benthamiana* at 22°C, fluorescence images were acquired using a confocal microscope (TCS SP5 or STELLARIS; Leica, Germany). GFP signals were excited at 488 nm and detected at 500–530 nm. For heat treatment, samples were incubated in liquid MS medium (0.5x MS mineral salts, 1% sucrose, 5 mM 2-(N-Morpholino) ethanesulfonic acid (MES; Sigma, Japan), pH adjusted to 5.7 with KOH) for 1 h at 37°C following the initial 36 h incubation.

For co-localization analyses, the full-length *DOT2* (*dot2^S^*and *dot2^C^*) coding region containing a stop codon was cloned downstream of the CaMV 35S promoter and fused in frame with mRFP in the p35S-FAST-mRFP vector. *Agrobacterium* cultures carrying *STA1 (sta1-1)-GFP* and *mRFP-DOT2* (*dot2^S^*and *dot2^C^*) constructs were co-infiltrated into *N. benthamiana* leaves and fluorescent signals were observed as described above. RFP signals were excited at 532 nm and detected at 580–620 nm.

For BiFC analyses, the same entry clones used for nuclear localization analyses were used. *STA1 (sta1-1)* and *DOT2* (*dot2^S^* and *dot2^C^*) entry clones were recombined downstream of the CaMV 35S promoter and fused in frame with nEYFP or cEYFP in the pSITE-nEYFP-N1 (GU734648) or pSITE-cEYFP-C1 (GU734652) Gateway vectors, respectively. *Agrobacterium* cells carrying *pSITE-STA1 (sta1-1)-nEYFP* or *pSITE-cEYFP-DOT2* (*dot2^S^*and *dot2^C^*) constructs were co-infiltrated in combination into *N. benthamiana* leaves in the indicated combinations, and fluorescent signals were observed as described above. EYFP signals were excited at 488 nm and detected at 500–530 nm. As a nuclear speckle marker, the full-length coding region of *U1-70K* (*U1 SMALL NUCLEAR RIBONUCLEOPROTEIN-70K*, At3g50670) was amplified and cloned into p35S-FAST-mRFP between the *Bam*HI and *Sal*I (Enzynomics, Korea) sites. As a nucleolus marker, *FIB2* (*FIBRILLARIN* 2, At4g25630)*-mRFP*, provided by Dr. Hyun-Sook Pai, was used (Park et al., 2021). All primers used to generate constructs are listed in Supplemental Table 3.

### Yeast two-hybrid assay

The full-length coding region of *STA1* or *sta1-1* was inserted into the pGADT7 vector, and *DOT2* or *dot2^S^* was inserted into the pGBKT7 vector. The resulting constructs, *pGADT7-STA1* (*sta1-1*) as prey and *pGBKT7-DOT2* (*dot2^S^*) as bait, were co-transformed into the *Saccharomyces cerevisiae* strain AH109 (Takara, Japan) and incubated at 30°C for 3 days. Transformants were selected on SD/-Leu/-Trp or SD/-Leu/-Trp/-His medium supplemented with 3-amino-1,2,4-triazole (3-AT; Sigma, Japan), a competitive inhibitor of the *HIS3* gene product. All primers used to generate constructs are listed in Supplemental Table 3.

### Arabidopsis mesophyll protoplast transient expression assay

Protoplast isolation and transient expression assays were performed as previously described (Kim *et al*., 2017). Effector constructs (*STA1*, *sta1-1*, *DOT2*, *dot2^S^*, and *dot2^C^*) and reporter constructs (g*HSFA3* fused to NanoLuc luciferase (nLUC)) were generated by inserting the corresponding coding sequences between the HBT promoter (a modified CaMV 35S promoter) and the NOS terminator in a plant expression vector. Firefly luciferase (fLUC), driven by the UBQ10 promoter, was included as an internal control.

For nuclear localization analysis in protoplasts, *HBT-DOT2* (*dot2^S^*and *dot2^C^*) constructs were transfected to *Salk007933^STA1-GFP^*or *Salk007933^sta1-1-GFP^*. Fluorescence images were acquired using a confocal microscope (TCS SP5; Leica, Germany). GFP signals were excited at 488 nm and detected at 500–530 nm.

For functional splicing assays, *HBT-DOT2* (*dot2^S^* and *dot2^C^*) constructs were transfected to *Salk007933^sta1-1-GFP^* and reporter activities were calculated as the nLUC/fLUC ratio. Transfected protoplasts were lysed in passive lysis buffer (Promega, USA) containing 1% Triton X-100 (USB, USA) and briefly vortexed. Lysates were incubated at room temperature for 10 minutes and then centrifuged at 13,000 rpm for 30 s. The supernatant was transferred for measurement of fLUC and nLUC activities using the Nano-Glo® Dual-Luciferase® Reporter Assay System (Promega, USA) and a GloMax 20/20 Luminometer (Promega, USA). All primers used to generate constructs are listed in Supplemental Table 3.

### Pollen viability

To assess pollen viability, 2-month-old plants were subjected to heat treatment (37°C for 1 day) or maintained under control conditions. Pollen grains were stained using the Alexander staining method as previously described (Peterson, 2010). Flower buds prior to anthesis were collected and fixed in Carnoy’s fixative (6 ethanol : 3 chloroform : 1 acetic acid) for at least 2 h. Fixed flower buds were then stained in the final staining solution at 50°C for 2 min. Stained pollen grains were observed using a light microscope (Leica ICC50, Germany).

### Propidium iodide staining

Seedlings from 3-day-old plants were collected and incubated in a 10 μg/mL propidium iodide (PI) solution (Sigma, Japan) for 1 min. Seedlings were then washed with distilled water and observed using a Leica TCS SP5 laser scanning confocal microscope (Leica, Germany). PI fluorescence was excited at 532 nm and detected at 590–630 nm.

### RNA-seq analysis

Total RNA was extracted from 7-day-old WT, *sta1-1*, and *S307* seedlings using RNAiso Plus (Takara, Japan). Heat stress (37°C for 3 h) was applied to 7-day-old plants, and heat-treated samples were collected. Three independent biological replicates were prepared for each condition. Sequencing libraries were constructed using the TruSeq Stranded mRNA Library Prep Kit with indexed adapters and RNA sequencing was performed using the Illumina NovaSeq platform (MacroGen, Korea). Paired-end raw sequencing reads were pre-processed to trim adaptors and low-quality reads using CLC Genomics Workbench v20.0.4 with default parameters (quality scores, 0.05; maximum ambiguous nucleotides, 2). Clean reads were mapped to *Arabidopsis* reference genome (TAIR10). Intron retention ratios at the whole-transcript level were estimated using IRFinder (v.1.3.0) (Middleton et al., 2017). Intron retention events were defined as those showing a change in IR ratio (ΔIR ratio) greater than 0.1 with an adjusted *p*-value (*p*_adj_) < 0.05. IR ratios were transformed into z-scores and visualized as heatmaps. Volcano plots were used to assess the overall distribution of IR changes in each comparison. RNA sequencing data have been deposited in the Gene Expression Omnibus (GEO) under accession number GSE316463.

## ACKNOWLEDGMENTS

We thank our lab members, as well as Dr. Inki Kim (Asan Medical Center, Korea) and Dr. Hyun-Sook Pai (Yonsei University, Korea), for technical assistance and materials. The authors declare no conflict of interest.

## AUTHOR CONTRIBUTIONS

H.K., K.-j.Y., S.P., and D.S. conducted the experiments and performed bioinformatic analyses. J.-H.J., D.-H.J., W.T.K., D.J.Y., and B.-h.L. advised on the research. All authors discussed the results. H.K. S.P., D.-H.J., and B.-h.L. wrote the manuscript.

---

**Supplemental Figure 1. Positional cloning and the identification of the *dot2^S^* mutation in *S307***

(A) Map of the *DOT2* locus on chromosome 5. Ninety-four F2 seedlings derived from an *S307* x L*er* cross were used for mapping. The Simple sequence length polymorphism (SSLP) markers used for mapping, Ch5_5.46Mb and Ch5_5.71Mb, are shown on the 184A1 and 35B3 BAC clones, respectively. Recombination numbers are shown under each marker locus.

(B) The mutation information of *dot2^S^* mutation, including nucleotide and corresponding amino acid changes.

**Supplemental Figure 2. Homologs and expression patterns of *DOT2*, a U4/U6·U5 tri-snRNP associated protein**

(A) Multiple sequence alignments of DOT2 homologs.

(B) Expression patterns of *DOT2* and *STA1::GUS* in *Arabidopsis*. GUS signals are observed in whole seedling, flower, silique, and trichome from left to right.

**Supplemental Figure 3. Phenotypes of the *dot2* single mutant alleles**

(A) Growth phenotypes of 3-week-old WT, *sta1-1*, *S307*, *dot2^S^*, and *dot2^T^* (SAIL_775_F10) plants (scale bar = 5 cm).

(B) Splicing defects in marker genes across genotypes. Box diagrams show spliced and unspliced isoforms. *PP2A* is used as a loading control.

(C) Representative confocal images of PI-stained root tips of 3-day-old WT, *sta1-1*, *S307*, *dot2^S^*, and *dot2^T^* seedlings (scale bar = 25 μm). Arrowheads indicate dead cells, and white bars mark the division zone. Division zone length in 3-day-old seedlings is quantified (n = 10). Statistical analysis is performed using one-way ANOVA with Tukey’s HSD test (\*\**P* < 0.01; n.s., not significant).

**Supplemental Figure 4. Residue positions of mutations in sta1-1 and dot2 mutant proteins**

(A) Predicted protein structures of *Arabidopsis* STA1 and DOT2. In the model, STA1 protein is shown in green and DOT2 protein in blue, with their respective mutations highlighted as follows: the *sta1-1* mutation in magenta, the *dot2^S^* mutation in red, and the *dot2^C^* mutation in orange.

(B) Domain structure of STA1 and DOT2. Predicted STA1-DOT2 interaction regions identified using InterProSurf and PDBePISA are indicated by red lines (STA1: 113–123 aa, 185–202 aa; DOT2: 548–569 aa). Domains are labeled as follows: UBQ, ubiquitin-like; PRP1P, pre–mRNA processing protein 1; HAT, half-A-tetratricopeptide repeat; NLS, nuclear localization signal; TPR, tetratricopeptide repeat; HIND, hub1–interacting domain.

(C) The *dot2^S^* and *dot2^C^* mutations, showing nucleotides and the corresponding amino acid changes.

(D) Growth phenotypes of 3-week-old WT, *sta1-1*, *S307*, and the *sta1-1 dot2^C^* double mutant (*ssd^C^d^C^*) plants (scale bar = 5 cm).

(E) Splicing defects in the marker genes across genotypes. The box diagrams show spliced and unspliced isoforms. *PP2A* is used as a loading control.

**Supplemental Figure 5. Rescue of *Salk007933* lethality through introduction of STA1-GFP or sta1-1-GFP, resulting wild-type- and *sta1-1*-like phenotypes, respectively**

**Growth phenotypes of 3-week-old *Salk007933^STA1-GFP^* and *Salk007933^sta1-1-GFP^* plants (scale bar = 5 cm).**

**Supplemental Figure 6. Increased STA1-aggregates under heat conditions**

(A and B) Subcellular localization of GFP-tagged STA1 and EYFP signals from BiFC analysis of STA1-STA1 interaction under heat conditions (scale bar = 75 μm).

(C and D) Co-localization of EYFP signals from BiFC analysis of STA1-STA1 interaction with the mRFP-U1-70K in leaves of *N. benthamiana* (scale bar = 10 μm).

(E) MolPhase prediction analysis of IDR region within STA1 (Liang *et al*., 2024).

For heat treatment, samples were incubated at 37°C for 1 h in liquid MS medium following a 36 h post-infiltration incubation.

**Supplemental Figure 7. Intron-retained *HSFA1* transcripts in *sta1-1***

(A) Increased IR events of *HSFA1s* in *sta1-1* (H). IR events in *WT*(H), *sta1-1*(H), and *S307*(H) are visualized using the Integrative Genomics Viewer (IGV).

(B) Protein domain structures of premature termination codon (PTC)-containing intron-retained pre-mRNA isoforms in *sta1-1* (H). Domains are labeled as follows: DB, DNA-binding domain; OD, oligomerization domain; NLS, nuclear localization signal; AHA1/2, aromatic and large hydrophobic amino acid residues embedded in acidic surroundings. PTCs are labeled with a red symbol.

Supplemental Figure 8. Splicing defects of *HSFA1*s in *sta1-1 dot2^C^* double mutants Splicing defects of the master heat stress regulators *HSFA1*s in the *sta1-1 dot2^C^* double mutant (*ssd^C^d^C^*) plants. Box diagrams show spliced and unspliced isoforms. *PP2A* is used as a loading control.

**Supplemental Table 1. Intron retained transcripts under normal conditions**

**Supplemental Table 2. Intron retained transcripts under heat conditions**

**Supplemental Table 3. Primer sets used in this study**

