## Supplementary figures and images for "The STA1–DOT2 interaction promotes nuclear speckle formation and splicing robustness in growth and heat stress responses"

### S307-SupplementalFigure1

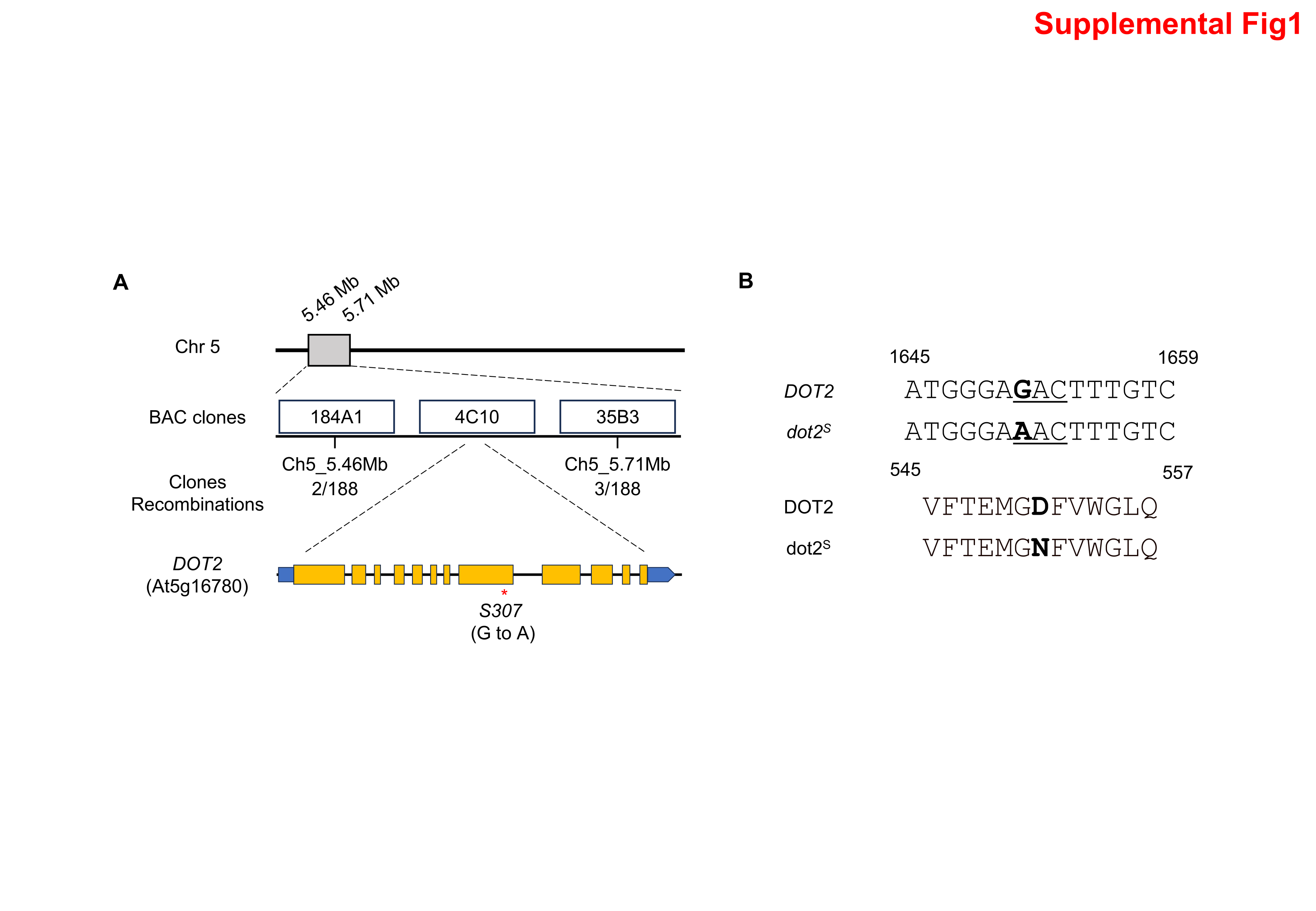

### S307-SupplementalFigure2

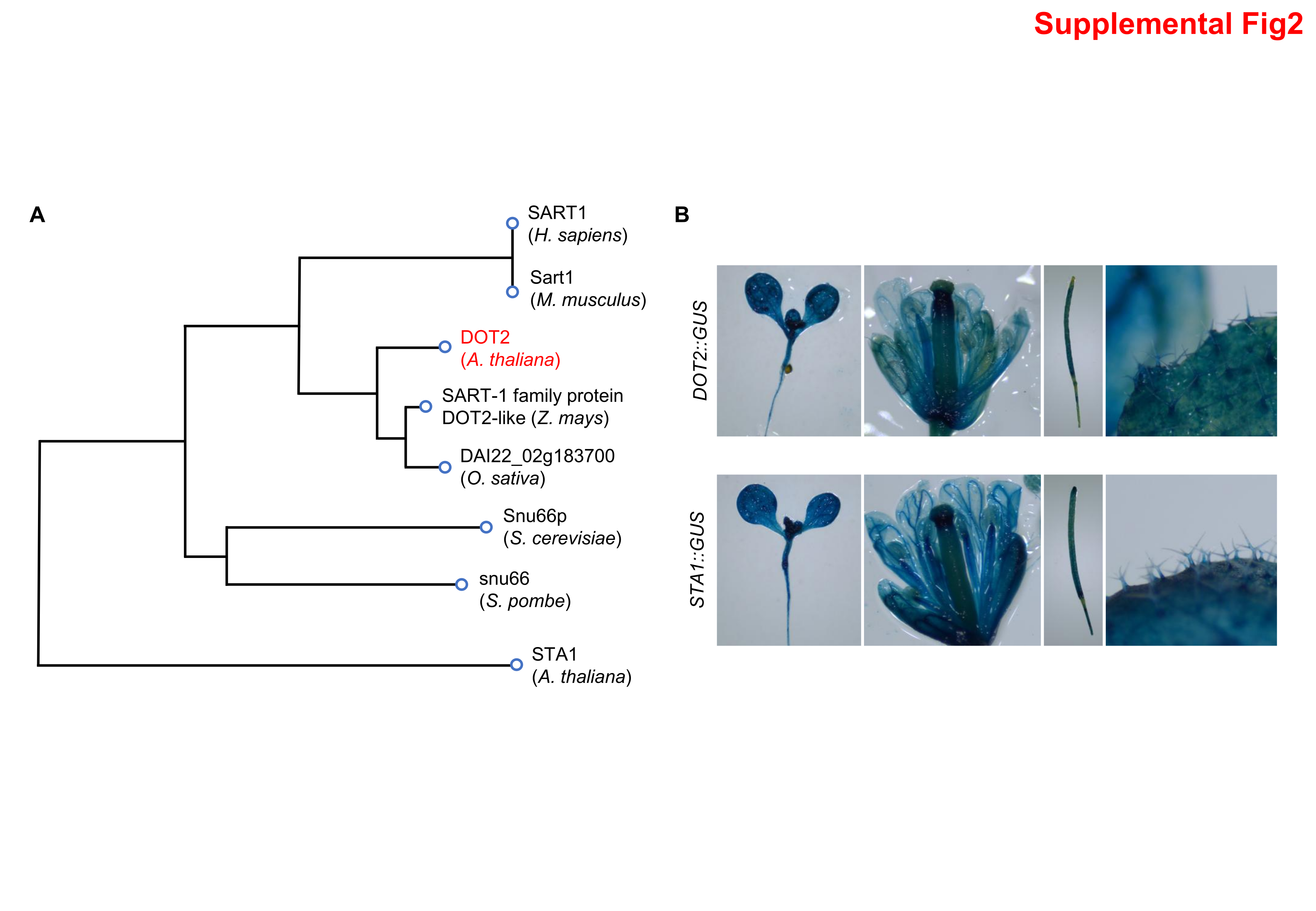

### S307-SupplementalFigure3

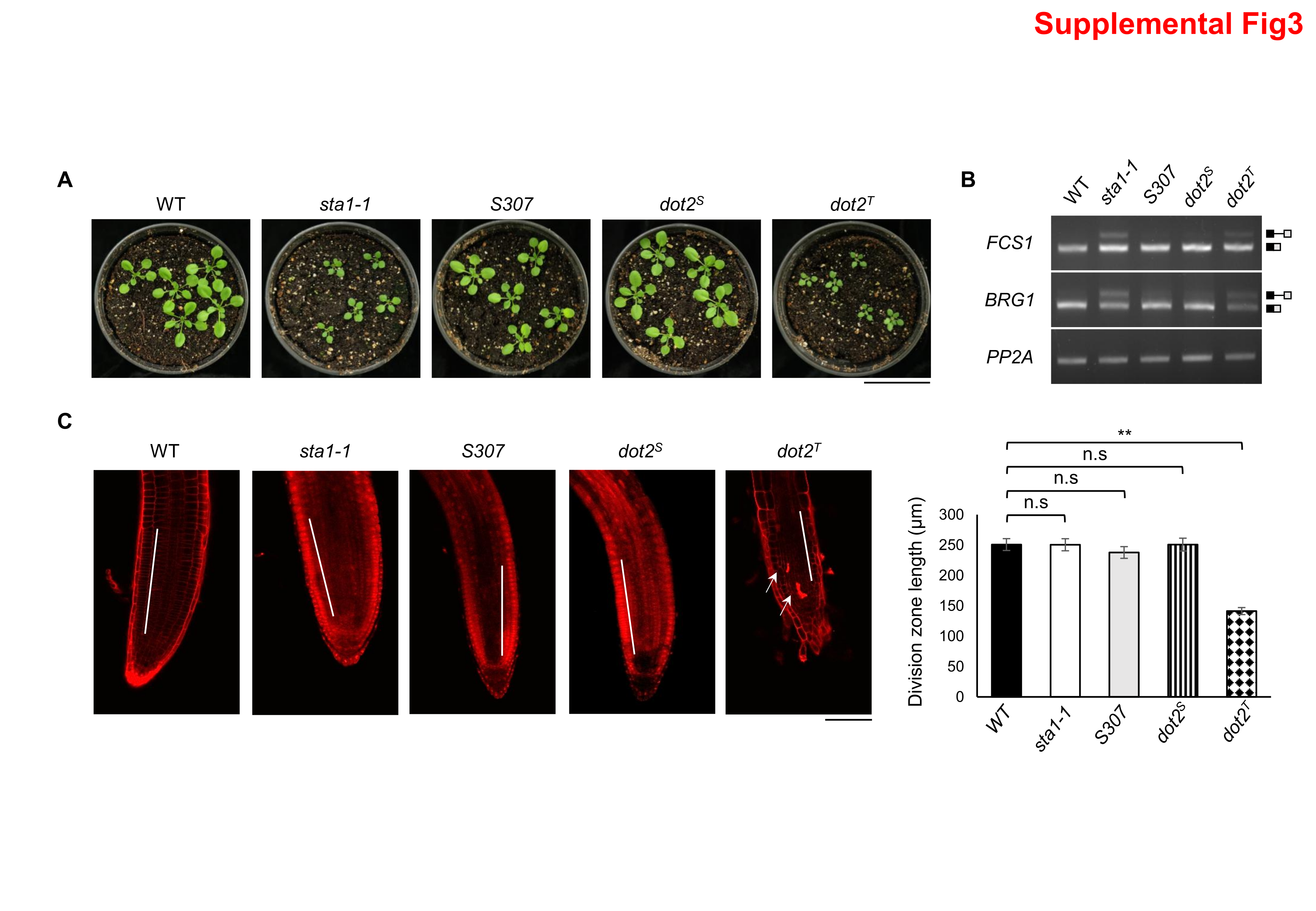

### S307-SupplementalFigure4

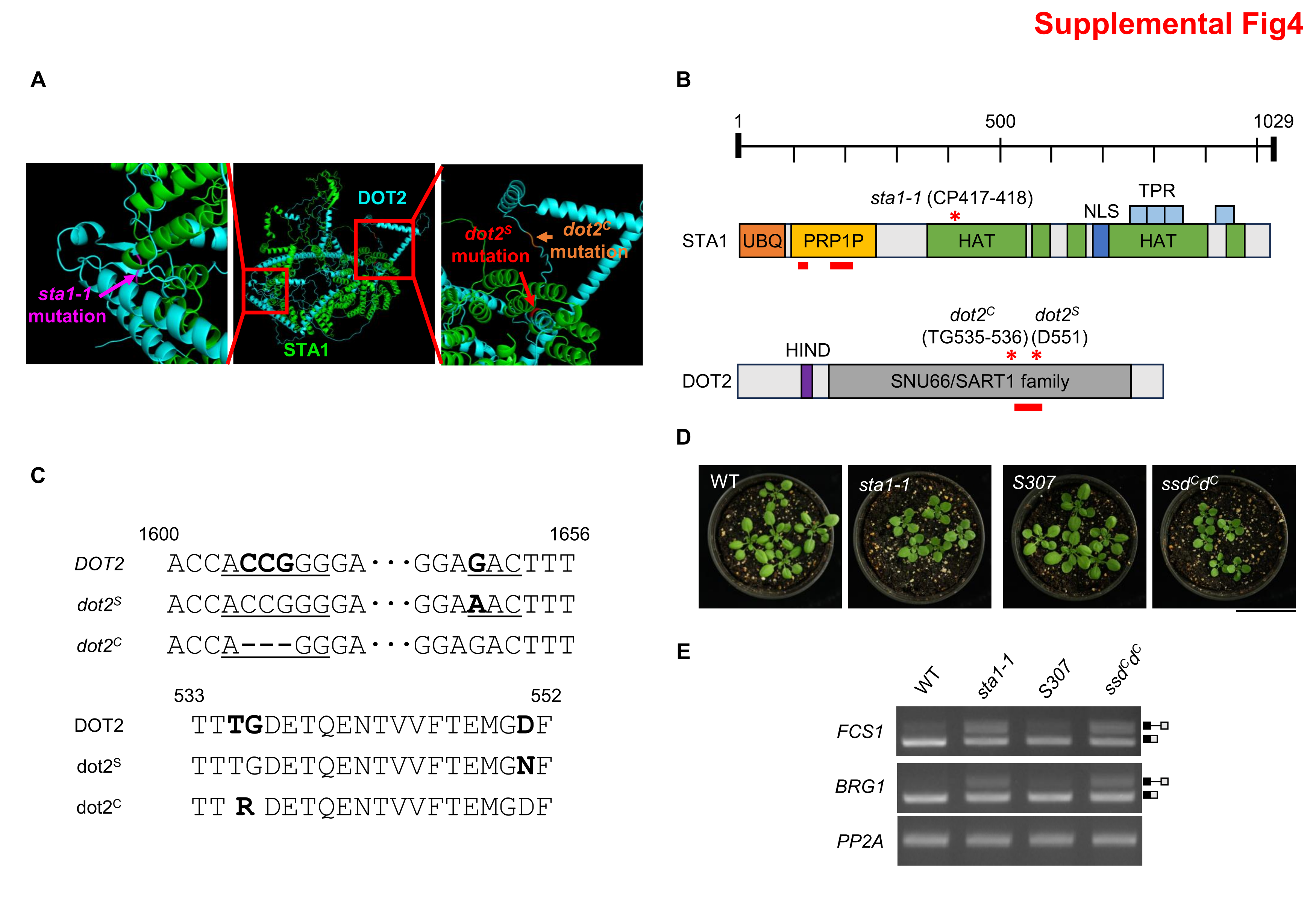

### S307-SupplementalFigure5

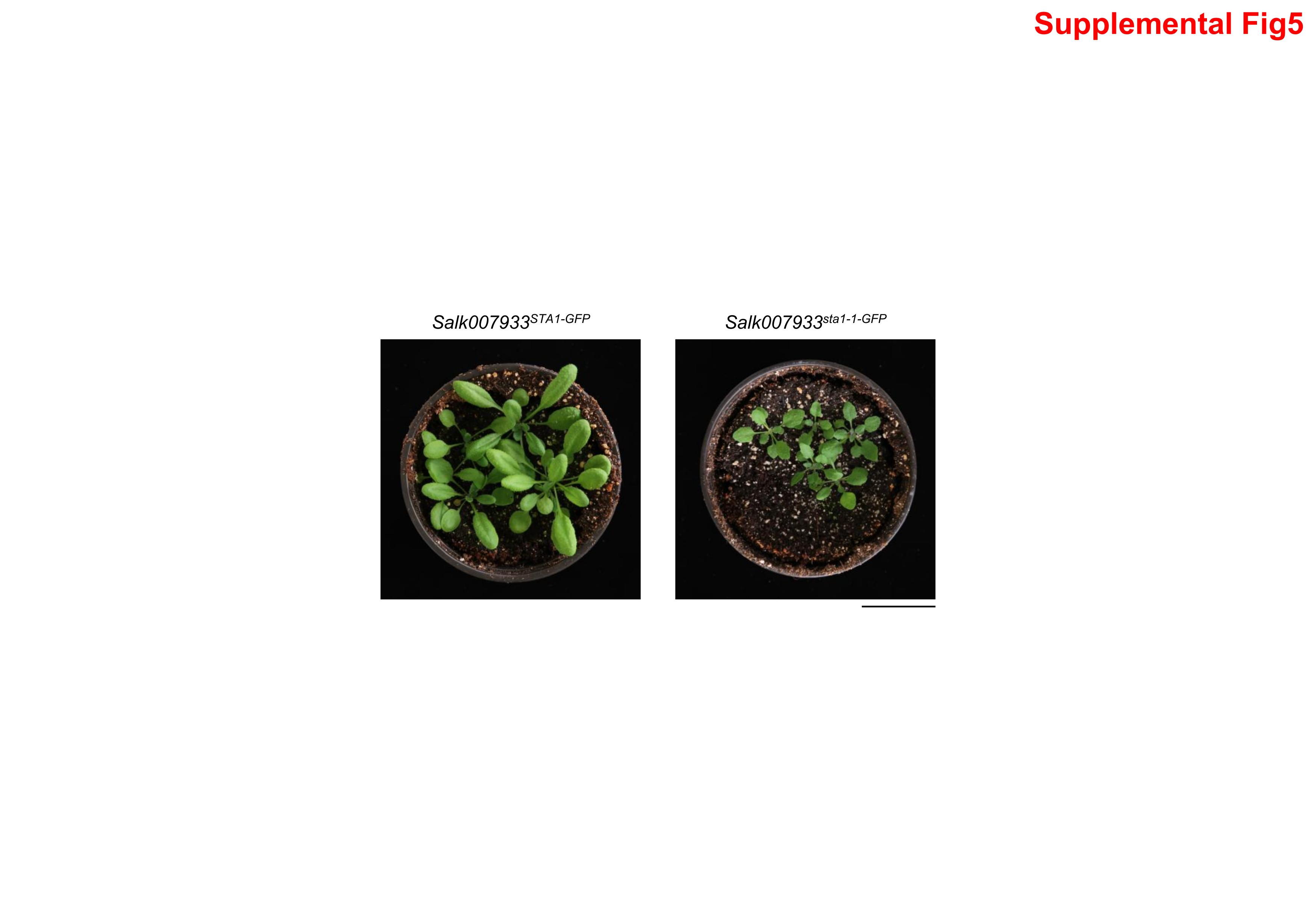

### S307-SupplementalFigure6

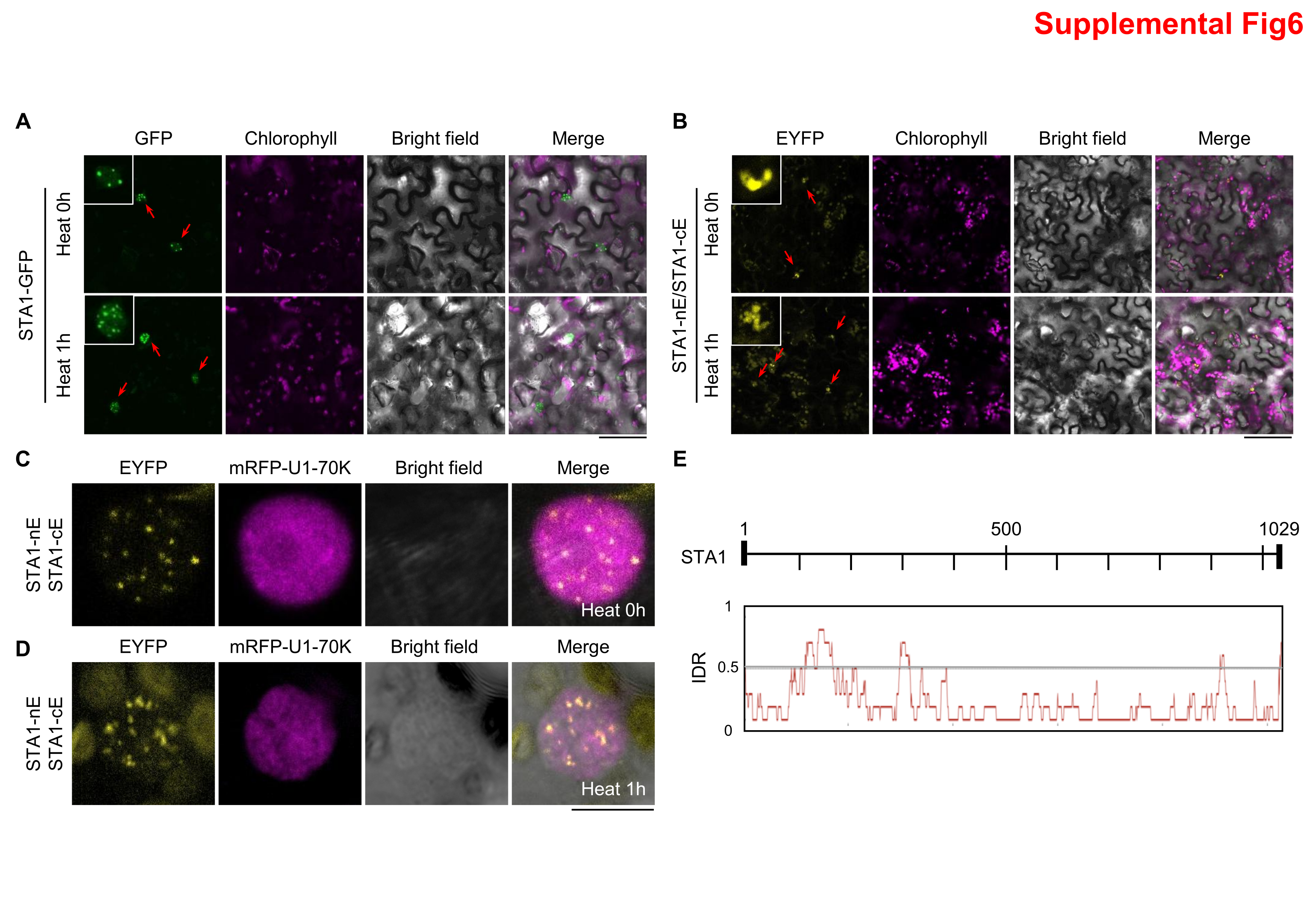

### S307-SupplementalFigure7

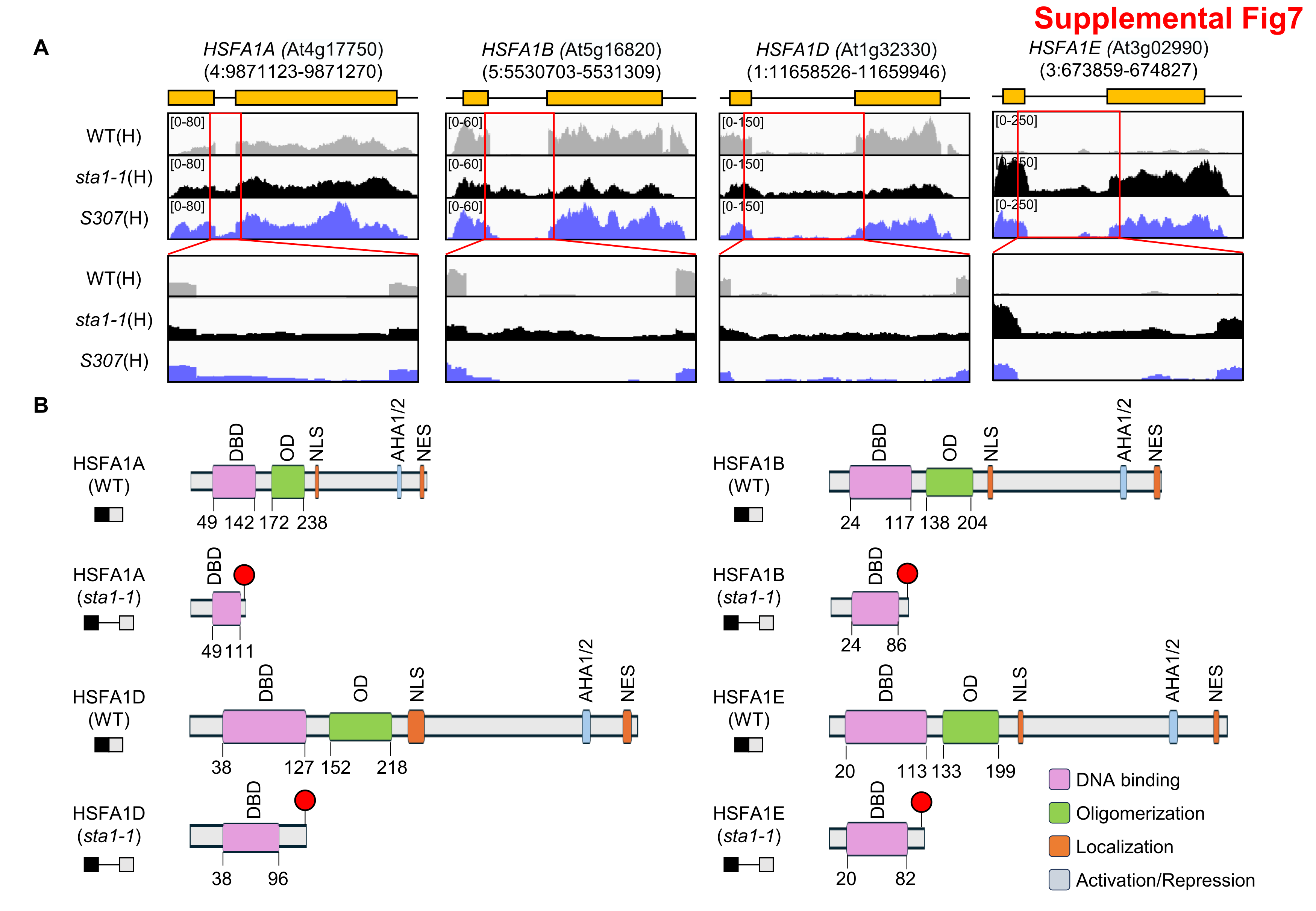

### S307-SupplementalFigure8

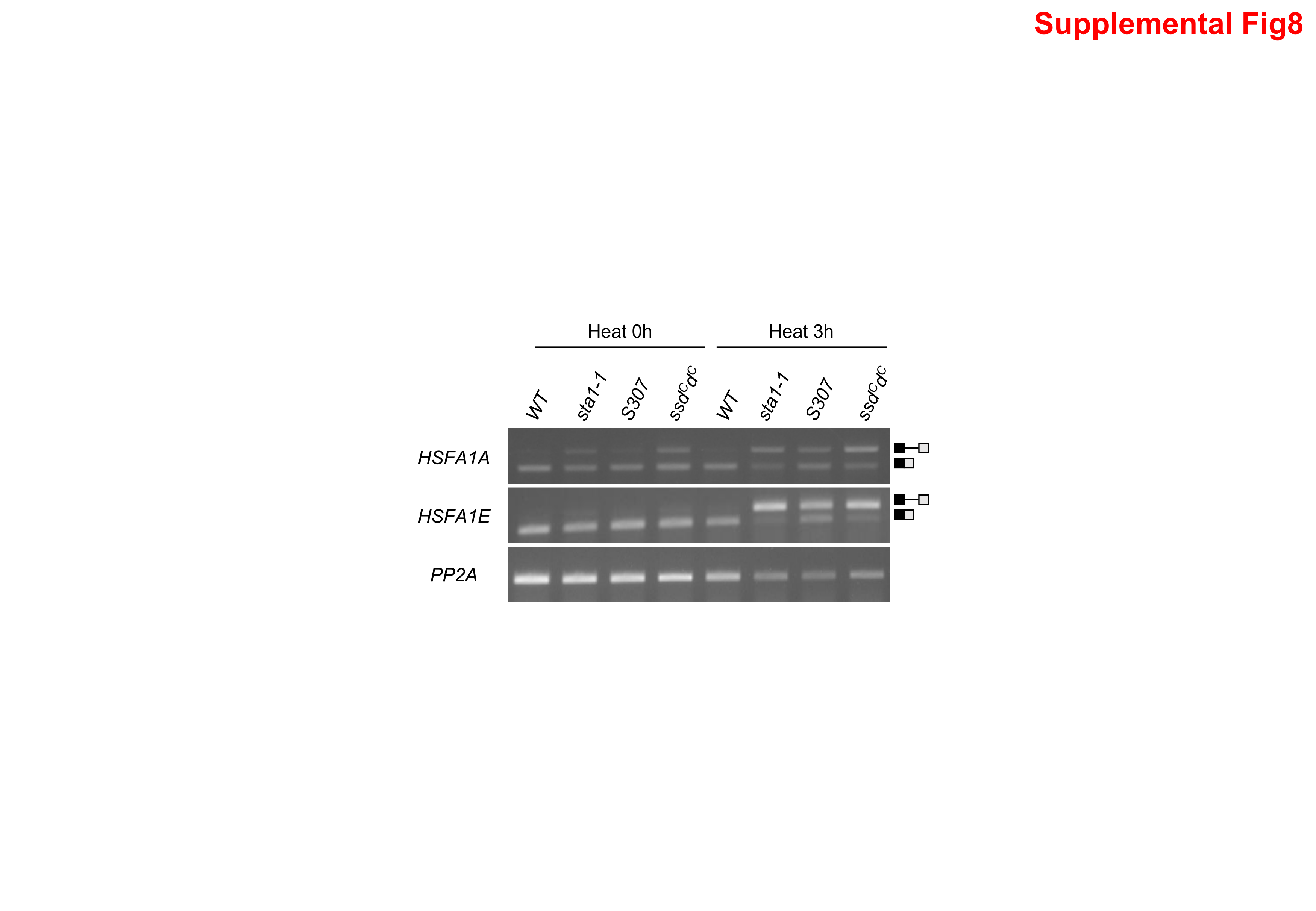
